# Microglial depletion worsens lesion in female but not male C57BL/6J mice after P10 hypoxia-ischemia

**DOI:** 10.1101/2023.10.17.562542

**Authors:** Danielle Guez-Barber, Sofia E. Nicolayevsky, Kaya J.D. Johnson, Sanghee Yun, Amelia J. Eisch

**Affiliations:** Division of Neurology, Department of Pediatrics, The Children’s Hospital of Philadelphia (CHOP), Philadelphia, PA, 19104, USA; University of Pennsylvania Perelman School of Medicine, Philadelphia, PA, 19104, USA; School of Arts and Sciences, University of Pennsylvania, Philadelphia, PA, 19104, USA; Department of Anesthesiology and Critical Care Medicine, CHOP Research Institute, Philadelphia, PA, 19104, USA; Department of Neuroscience, University of Pennsylvania Perelman School of Medicine, Philadelphia, PA, 19104, USA

**Author notes:** Author email addresses.

**Keywords:** microglia, hypoxic-ischemic encephalopathy, neonate, sex differences, macrophage, hippocampus

## Abstract

**Background:** Rodent models for perinatal hypoxic ischemic (HI) encephalopathy have reported sex differences such as the same injury causing larger lesions in the brains of males compared to females. Microglia, the resident immune cells of the brain that have sex-dependent developmental trajectories and gene expression patterns, likely play a different role in females and males following HI. However, there is conflicting literature on whether depletion of microglia worsens or improves HI-induced lesions and whether this differs by sex. Here we tested the effect of pharmacologic microglia depletion on HI lesion size in male and female mice.

**Methods:** An initial cohort of C57BL/6J mouse pups underwent HI at postnatal day 10 (P10) using a modified Vannucci procedure or a Sham insult followed by brain collection at P13. Another cohort of mice received daily intraperitoneal injections from P7 to P12 of either 25mg/kg PLX3397 (PLX, a CSF1R inhibitor) or vehicle (Veh). These mice also underwent HI or Sham at P10, resulting in four groups (Veh-Sham, Veh-HI, PLX-Sham, PLX-HI). All groups included female and male mice. Behavioral testing was performed both pre-HI (forelimb grasping [P8, P9]) and post-HI (open field traversal [P12], behavior and appearance observations [P13]). P13 brain sections underwent immunohistochemistry for Iba1 or cresyl violet staining for lesion scoring.

**Results:** P13 HI hippocampal sections had more Iba1 signal than Sham, with more variance in Male-HI vs Female-HI mice. PLX led to >95% depletion of Iba1+ cells at P10 or P13, and effective elimination of microglia did not differ by sex. In the hippocampus, Female-PLX-HI mice had worse lesion scores than Female-Veh-HI mice; this was not true in male mice, where there was a trend in the opposite direction. Female-PLX-HI mice also had worse lesion scores than Male-PLX-HI mice. In contrast to this sex-dependent effect of PLX on lesion score, there was no difference among groups in developmental milestones.

**Conclusion:** PLX3397 injection P7-P9 or P7-P12 effectively depletes microglia by P10 or P13, respectively. Microglial depletion via PLX worsens HI-induced injury in female mice but not in male mice.

## 1. Introduction

Perinatal hypoxic ischemic encephalopathy (HIE) is among the most common causes of death and disability in neonates globally.^1–3^ Multiple studies in perinatal HIE animal models report distinct outcomes by sex. For example, male mice have larger lesions than female mice after the same injury and perform worse than females on a hippocampal-dependent task.^4^ Several studies have shown that male rodents have larger anatomic injuries^4–6^ and worse cognition^7,8^ after hypoxia-ischemia (HI) relative to female rodents. The sex-specific differences are not limited to cognitive effects. Rats undergoing a developmental hypoxic injury show sex-dependent motor outcomes; males present a more spastic phenotype and females present a more dystonic phenotype.^9^ Even treatment approaches need to be assessed for their sex-specific effects. For example, while therapeutic hypothermia reduces HIE injury in humans and HI injury in rodents, meta-analysis of the effect in rats revealed that the beneficial effect of therapeutic hypothermia is driven by its effect in female, but not male, rats.^10^ Overall, these and many other studies underscore the critical importance of reporting and considering sex in HI studies.

Many factors may contribute to the sex-dependent response to perinatal HI, including sex differences in brain development^11–13^, cell death^14–16^, metabolism^17^ and immune function.^18–23^ One established factor mediating sex differences in stroke^24^ and other pathologies^25,26^ — but understudied in perinatal HI — is microglia. In healthy brains, microglia (immune cells native to the brain^27,28^) orchestrate developmental pruning of synapses^29–31^, shape cellular indices of learning^32,33^, and regulate behavior^34,35^ including sex-specific behavior.^25,27,36^ Microglia are different in males and females; they have sex-dependent developmental trajectories^34,37–39^ and gene expression patterns^40,41^. Lesion^42,43^ and transplantation^24^ studies show microglia play different roles in male and female brains after HI or stroke. However, there are conflicting data on whether microglial depletion prior to HI or stroke worsens lesion size^42,44,45^ or ameliorates injury^46^.

This discrepancy in the role of microglia in the context of HI may be in part explained by the method of microglial depletion. Pharmacologic depletion with CSF1R inhibitors can also impact peripheral monocyte-derived microglia-like macrophages^47,48^; genetic approaches with diphtheria toxin A cause a peripheral inflammatory response^48,49^ so may actually be stimulating peripheral macrophages. The role of sex in HI microglial depletion studies is unclear; studies were completed only in male mice^44,45^ or did not disaggregate data by sex.^46^ One study^42^ compared male and female mice and found microglial ablation aggravates injury only in males; the tamoxifen required to induce the genetic ablation of microglia may confound the reported sex differences. Overall, these conflicting data highlight a persistent knowledge gap: are microglia helpful or harmful in perinatal HI?

The present study had two goals. First, we aimed to assess the impact of postnatal day 10 (P10) HI or a control procedure (Sham) on pathology, including lesion size and microglia density, on male and female mice. We report here brain pathology after HI that demonstrates a subtle but clear sex difference. Second, we aimed to determine if prior microglial depletion exacerbated or ameliorated HI-induced pathology, and whether this varied by sex. Pharmacological depletion of microglia has traditionally been accomplished using CSF1R inhibitors, such as PLX3397, given orally over ∼2 weeks^50,51^. Such slow depletion of microglia is untenable for perinatal HI studies where the injury is induced at P10 and the brain examined at P13, and where depletion close to P0 may disrupt development prior to the P10 injury. To address this challenge, we initiated a PLX3397 injection regimen at P7 that largely depletes brain microglia by P10 compared to vehicle-injected (Veh) mice. We then employed this novel PLX3397 regimen to assess how microglial depletion impacts HI-induced pathology. Our data reveal a clear sex difference: in female HI mice, microglial depletion makes the lesion worse, while in male HI mice microglial depletion has a trend to decrease the lesion. These data are discussed in light of the complex role of microglia in health and disease.

## 2. MATERIALS AND METHODS

Additional details for each subsection in **Supplementary Materials and Methods (Supp. Methods)**.

### 2.1. Animals

Dams and males were C57BL/6J mice shipped from Jackson Laboratory and bred in the Children’s Hospital of Philadelphia (CHOP) Research Institute. Mouse pups were kept in the same cage as the dam and their littermates following birth and were not disturbed for 6 days. All mice were cared for in compliance with Protocol #1245 approved by the Institutional Animal Care and Use Committee at CHOP and guidelines established by the NIH’s *Guide for the Care and Use of Laboratory Animals.* Our scientific reporting adheres to the ARRIVE 2.0 guidelines.^52^

### 2.2. PLX3397 injections

PLX3397 (PLX, Selleck, Catalog No.S7818) was resuspended in dimethyl sulfoxide (Cell Signaling Technology, Catalog No.12611P) to make a 50mM stock stored at -80°C. Mice received daily ∼9AM intraperitoneal injections (*ip*) of 20μL PLX or injections of the same volume 20μL Vehicle (50% phosphate-buffered saline PBS, 50% polyethylene glycol). Brains were collected 24 hours after the last injection. Injection quality was noted each day for each mouse, and most days in most mice there was no residue on abdomen after *ip* injection.

### 2.3. Experimental groups

Four cohorts of mice were included in this study. In each cohort, mice of each sex were randomly assigned to experimental groups evenly distributed within each litter. <u>Cohort 1</u> received no drug treatment and underwent HI or Sham at P10 for assessment of Iba1 response and presence of pale lesion at P13; this included 140 pups. <u>Cohort 2</u> was used for a PLX3397 dose-response study and included 18 pups across three experimental groups: Veh, 25mg/kg PLX3397, and 50mg/kg PLX3397. <u>Cohort 3</u> included one litter of 9 mice: 50mg/kg PLX3397 (n=5) or Veh (n=4). Of the mice receiving 50mg/kg PLX mice, 80% died by P12; there were no deaths in the Veh groups. <u>Cohort 4</u> included the variables of drug (25mg/kg PLX3397 or Veh) and injury (HI or Sham). This included 78 pups total: 38 females, 40 males.

### 2.4. Behavioral Testing

Developmental milestones and behavior were tested on postnatal days 8, 9, and 12 as previously published.^53^ At P8 and P9, forelimb grasp (strength) was measured; at P12, exploration was assessed.

### 2.5 Qualitative Appearance Assessment

At P10, fur state was assessed prior to HI; mice with pinker abdomens were noted. At P13, fur state, including hair loss, and gross behavioral observations of pups were noted when the dam was removed, first in the home cage for 5min, and then in a novel environment with littermates for an additional 5min.

### 2.6. P10 Hypoxia-Ischemia

A modified Vannucci Model^2,54–56^ of HI was used in male and female PLX and Veh mice at P10. Briefly, lidocaine ointment was placed on the pup’s ventral neck, then under isoflurane anesthesia (3% v/v for induction and 1% v/v for maintenance), a midline ventral incision in the anterior neck was made. HI pups underwent double ligation of the right common carotid artery with a 5.0 silk suture; Sham pups underwent surgery including incision and isolation of the right carotid artery without ligation. The entire surgery lasted ∼5-7min. After carotid ligation or Sham surgery, the mice recovered for 60-80min on a heating pad with their littermates; temperatures of each individual pup were closely monitored and maintained between 36.0-37.9°C. Recovery was followed by 45min of hypoxia exposure where oxygen concentration was decreased to 8% with balanced nitrogen in a controlled chamber (BioSpherix Ltd., Parish, NY, USA). The Sham mice remained the same amount of time in normoxia conditions (21% FiO_2_) in a similar chamber. Across all experimental litters, the total time pups spent away from their dam was on average 120min. Within each litter, time away from the dam was calculated for HI pups and averaged; all Sham mice spent exactly this time away from the dam.

### 2.7. P10 Seizure Scores

During hypoxia as well as 10min before and 10min after, mice were assessed on a modified Racine scale for seizure activity. Briefly, every 5min pups were assigned a score from 0 to 6 to reflect maximal seizure activity in that bin of time; cumulative seizure score was generated for each pup.

### 2.8. Brain Collection and Tissue Preparation

Mice were euthanized at P10 or P13 via rapid decapitation in compliance with NIH guidelines for euthanasia of rodent neonates. Brains were placed in room temperature (RT) 4% paraformaldehyde (PFA) and fixed for 72 hours. Following two PFA changes (1/day), brains were cryoprotected in 30% sucrose in 0.1 M phosphate-buffered saline (PBS) at 4°C. Forty µm coronal sections were collected on a freezing microtome (Leica SM 2000 R) and stored in 1xPBS with 0.01% sodium azide at 4°C until processing for histological assessment.

### 2.9. Pale Surface Brain Lesion at P13: Presence and Size

For the presence of a pale lesion on the surface of the brain, each extracted brain underwent gross visual examination. If present, these pale brain surface lesions were always on the right hemisphere since they result from right carotid artery ligation. If a pale surface lesion was present, a flexible ruler was placed on the surface of the brain to measure (in 0.5mm increments) the lesion at its greatest extent rostro-caudally and medio-laterally; calculated lesion area was the product of the length x width of the lesion.

### 2.10 Immunohistochemistry (IHC) for Iba1

IHC was performed as previously described^37,57,58^ using a primary antibody against Iba1, rabbit anti-Iba1 (1:5000; Wako Chemicals, #019-19741), which labels both brain-resident microglia and monocyte-derived macrophages, and a biotinylated donkey-anti-rabbit IgG (1:200; Jackson ImmunoResearch Laboratories, #711-065-152) secondary antibody. Signal was amplified using avidin-biotin complex and visualized using diaminobenzidine.

### 2.11. Quantification of Iba1-immunoreactive (+) cells

Methods 2.11.1 through 2.11.3 used images from Iba1-immunostained (Iba1+) sections from P13 mice captured in bright-field light microscopy using an Olympus BX51 microscope with a 10x/0.3NA objective and an Olympus DP74 camera (widescreen aspect ratio, 16:10). The approach to quantify Iba1+ cells is described below.

#### 2.11.1. Iba1+ pixel percentage in ipsilateral hippocampus after Sham or HI (Cohort 1)

For every mouse in Cohort 1, four 100x images of the ipsilateral (right) hippocampus per brain were obtained at -1.70, -2.30, -2.92mm relative to bregma, with two images at -2.92mm to capture the entire hippocampus at that bregma. Images were analyzed in Fiji to obtain the percentage of pixels positive for Iba1 immunoreactivity. These were averaged across the four images per brain to obtain a single value representing the percent of Iba1+ pixels for the hippocampus of each mouse.

#### 2.11.2. Iba1+ cell density in left striatum, cortex, and hippocampus after Veh or PLX (Cohort 2)

For every mouse in Cohort 2, eight 100x images per brain were taken from five left hemisphere sections ranging from +1.33 to -2.91mm from bregma; these sections included cortex, hippocampus, and striatum. Guidelines to ensure similar brain region representation and calculation of Iba1+ cell density are described in **Supp. Methods.** Iba1+ cell density was collected for each brain region (striatum, cortex, hippocampus) present in each photomicrograph. Values were averaged so that each mouse had one value for the striatum, cortex, and hippocampus. Iba1+ cell densities from these regions were averaged such that each mouse was represented by one density value.

#### 2.11.3. Iba1+ cells in contralateral cortex after Veh or PLX and after Sham or HI (Cohort 4)

For every mouse in Cohort 4, one 100x image per brain was obtained at +1.21mm from bregma, maximizing the area within the image that contained the contralateral (left) cortex. In Fiji, the cortex was outlined to exclude subcortical regions. This area of the cortex was measured and converted to mm^2^. The number of Iba1+ cells in this image of cortex was counted for both Veh and PLX mice; however, they were exhaustively counted throughout the cortex in this image in PLX mice, and counted within a 400×400 pixel square in Veh mice. The density of Iba1+ cells in the cortex was calculated.

#### 2.11.4. Iba1+ cells in ipsilateral and contralateral hemisphere in PLX mice after Sham or HI (Cohort 4)

For every PLX mouse in Cohort 4, three sections (+1.21mm, -2.27mm, and -3.63mm from bregma) were scanned at 20x using a brightfield Aperio AT2 digital slide scanner from Leica Biosystems. Iba1+ cells were quantified using QuPath^59^. The area of each hemisphere (in mm^2^) was measured and the Iba1+ cells were manually counted throughout each hemisphere, resulting in the number of Iba1+ cells per mm^2^ in each hemisphere.

### 2.12. Cresyl Violet Staining and Lesion Scoring (Cohort 4)

Forty µm coronal brain sections in a 1:12 sampling ratio ranging from +1.41 to -3.63mm relative to bregma were stained with 0.1% cresyl violet acetate (CV). Sections were analyzed in bright-field light microscopy. Lesion scoring for every section (range 7-11 sections per brain) was performed by two independent observers masked to experimental conditions in four hippocampal subregions (CA1, CA2, CA3, dentate gyrus), striatum, and cortex. The left and right hemispheres were each scored independently. The entirety of each region within a section was scored on a scale of 0-3 with the following criteria: 0=no injury, 1=scattered shrunken neurons, 2=moderate cell loss with infarction in columnar distribution, 3=severe cell loss with cystic infarction.^60^ For each brain section, a score of 0-3 was given to each brain region analyzed. For each brain within each subregion, scores across bregma were averaged to yield average subregion lesion scores. Average subregion lesion scores and an average ipsilateral lesion score (average of all 6 subregions) from the two observers were averaged together to yield the final score graphed and analyzed; the average difference in lesion scores between observers was 1.6%. To visualize lesion severity across groups in another way, the fraction of mice with average lesion score within each 0.5 increment bin was visually represented as parts of a whole.

### 2.13. Statistical Analysis

All surviving mice were included in all statistical analyses except one Female-PLX-HI mouse. This mouse was an outlier in Iba1 cell counts (Grubbs test, G=3.331, *p<0.05) and its data were removed throughout; this mouse also had notable PLX residue on its abdomen after PLX injections on both P9 and P10. These exclusion parameters were not determined *a priori*. Additional details, including statistical testing approach, are included in **Supp. Methods.**

**Table 1**. This table includes the sample size, statistical tests and results for all Figures included in the main text. See **Supp. Methods** for statistical details, and Supp. Table 1 for sample size, statistical tests and results for Supp. Figures.

**Table 1.**
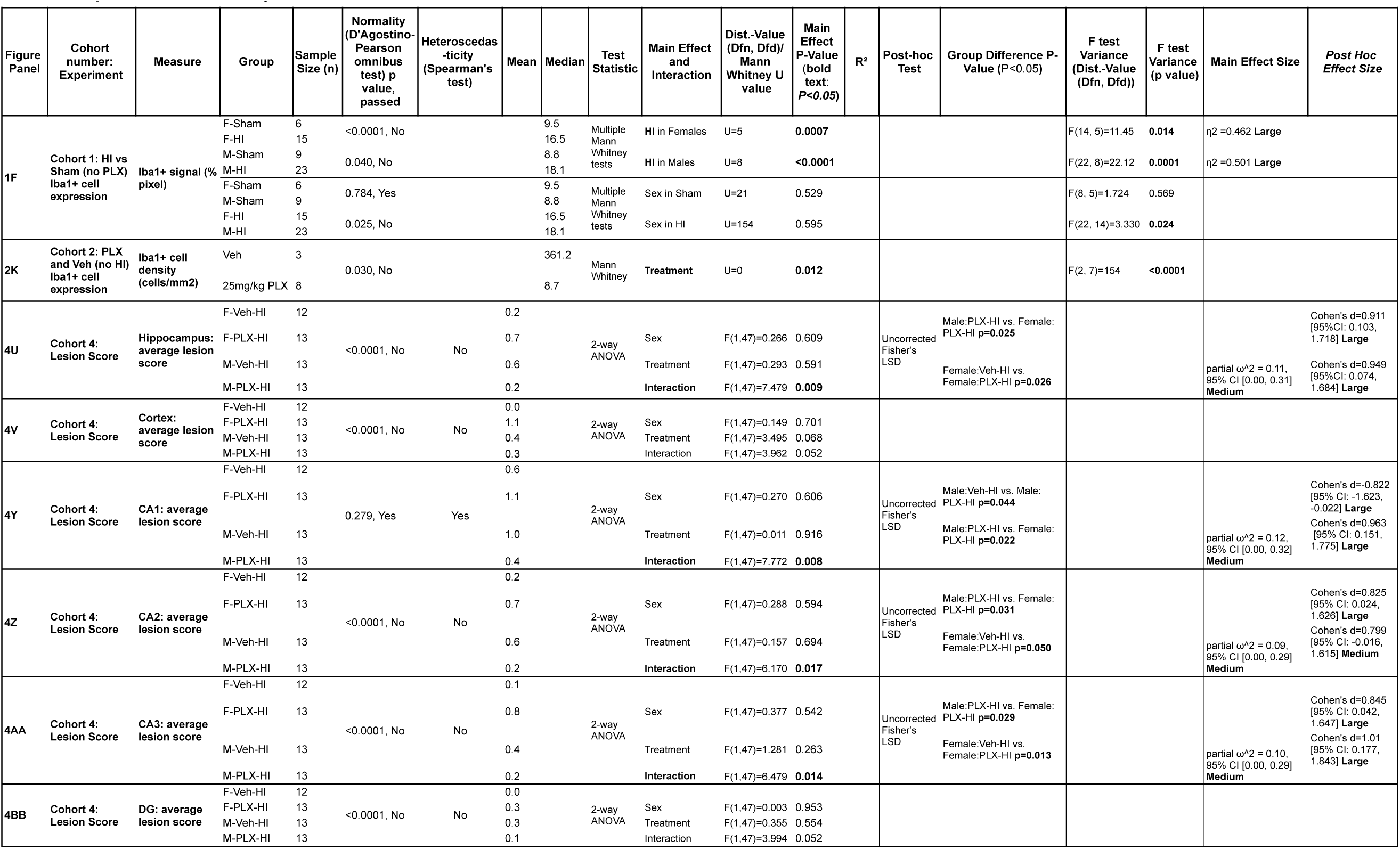
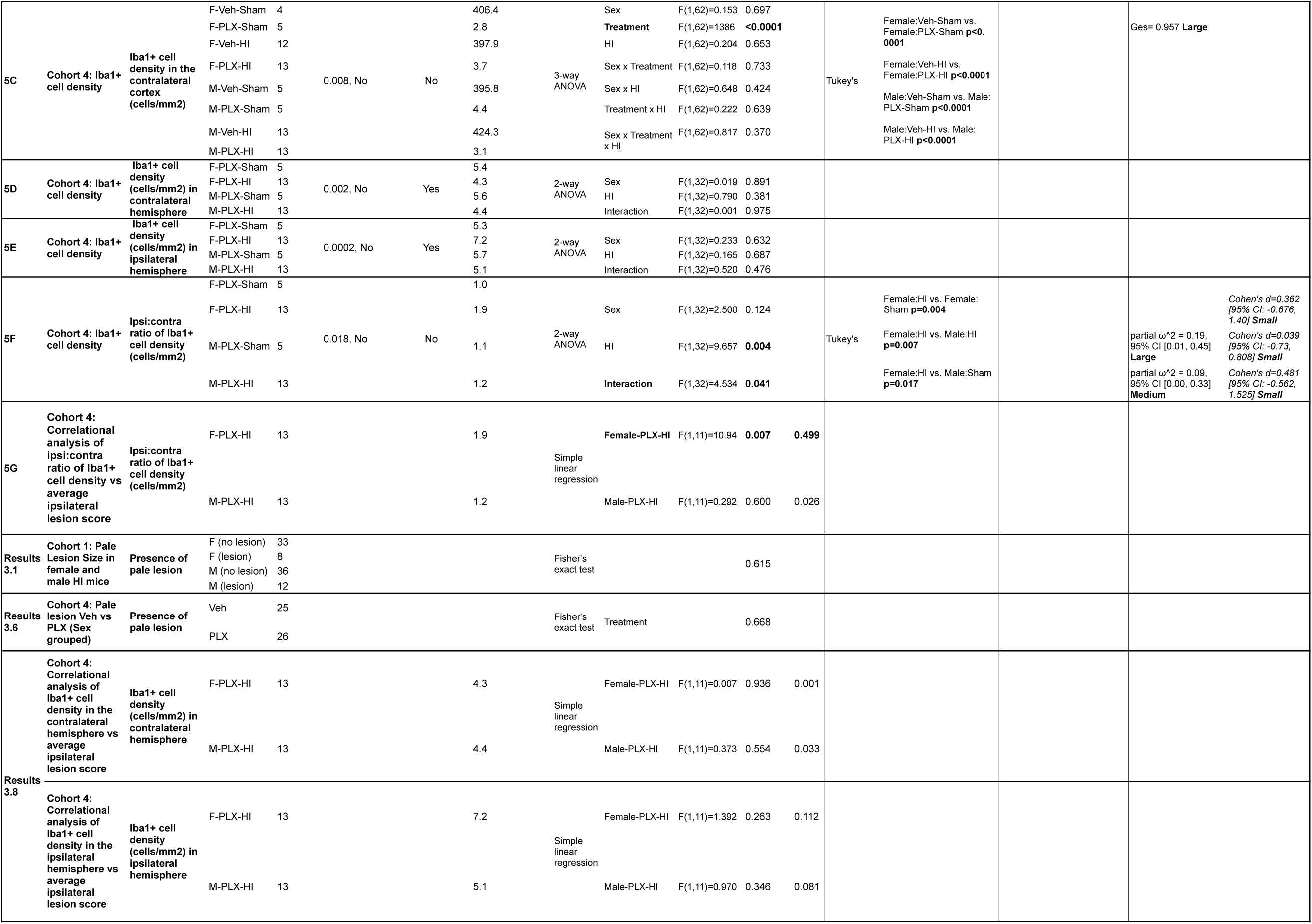
Sample Size, statistical analysis and results.

## 3. RESULTS

### 3.1. HI induces a pale lesion on the brain surface which is larger in Male-HI vs Female-HI mice

Mice underwent Sham or HI at P10 and brains were removed at P13 **(Fig. 1A)**. In Cohort 1, whole brains were first examined for gross pathology, specifically a pale lesion evident on the surface of the brain **(Supplementary [Supp.] Fig. 1)**. No Sham mice had a pale lesion on the brain surface, as expected (Sham n=45 in Cohort 1). Of all 89 HI mice in Cohort 1, 22% (20/89) had a detectable pale lesion on the surface of the brain **(Table 1)**. When considered by sex, 20% of the Female-HI mice (8/41) and 25% of the Male-HI mice (12/48) had a pale brain surface lesion **(Table 1)**. While there was no sex difference in the presence of a pale lesion **(Table 1)**, when considering only the mice who had a pale surface lesion, the lesion was >2-fold larger in Male-HI mice vs Female-HI mice, and there was more variance of lesion size in Male-HI compared with Female-HI mice **(Supp.Fig. 1)**. These data show that in HI mice that show a pale lesion on the brain surface, the lesion size is larger in male vs female mice.

**Figure 1.**
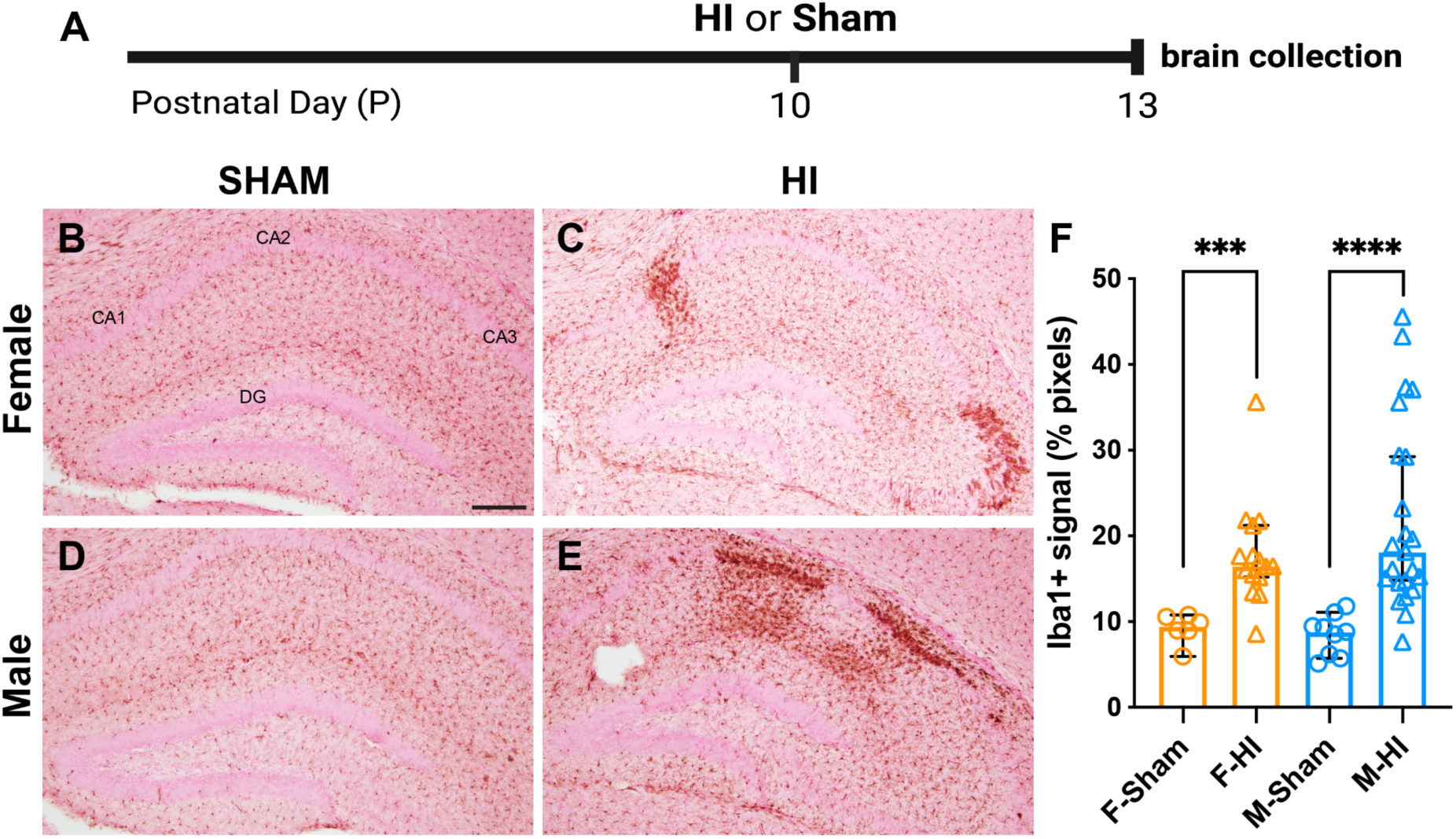
Iba1 is increased by hypoxia ischemia and levels vary more in Male-HI vs Female-HI mice. **(A)** <u>Timeline for Cohort 1</u>: HI or Sham was induced at P10, and brains were collected at P13. **(B-E)** <u>Representative images</u> of anterior hippocampal sections from P13 mice including Female-Sham mouse **(B)**, Female-HI mouse **(C),** Male-Sham mouse **(D)**, and Male-HI mouse **(E)**. Iba1-immunoreactive (+) cells were visualized, likely reflecting both brain-resident microglia and monocyte-derived macrophages that infiltrate the brain from the peripheral blood after injury. Subregions of the hippocampus indicated: CA1, CA2, CA3, dentate gyrus (DG). **(F)** <u>Percent of Iba1+</u> <u>pixels</u> in the hippocampus of P13 Sham and HI Cohort 1 mice. HI mice had higher percentages of Iba1+ pixels than Sham mice. There was more variance of Iba1+ signal in HI vs Sham in both Female and Male mice, and more variance in Male-HI vs Female-HI. Magnification in **(B-E)** 100x; scale bar in **B**, 200µm. F, Female. HI, hypoxia ischemia. M, Male. Complete details in **(Table 1).**

### 3.2. HI induces Iba1 in Female and Male mice, with greater variation in Male-HI vs Female-HI mice

Iba1 is a protein expressed in the brain by both microglia and infiltrating monocyte-derived macrophages, and an increased number or density of Iba1+ cells is often seen after HI.^5,61,62^ To assess Iba1 after HI and potential sex-dependent HI effects on Iba1, Cohort 1 mice underwent P10 HI or Sham and at P13 their brains were collected for examination of Iba1+ cells **(Fig. 1A)**. Qualitatively, Iba1+ cells were evident in the hippocampi of all Sham and HI mice **(Fig. 1B-E)**. In Sham mouse brains, Iba1+ cells were evenly-spaced within recognizable brain regions and hippocampal subregions **(Fig. 1B, D)**. In P13 Sham mouse brains, there was no qualitative difference in Iba1+ hippocampal cells in Female vs Male mice, in keeping with prior work from P10 mice.^37^ In contrast, in P13 HI mice, Iba1+ cells were irregularly-spaced, with dark patches of cells appearing in an unpredictable pattern **(Fig. 1C, E)**. Quantitative assessment of Iba1+ pixels in all four groups of mice revealed Female-HI mice had 194% more vs Female-Sham mice, and Male-HI mice had 261% more vs Male-Sham mice **(Fig. 1F)**. There were also HI- and sex-dependent differences in variance. HI values varied more than Sham values within each sex. While Female-Sham and Male-Sham values had similar variance, Male-HI values varied more than Female-HI values **(Fig. 1F)**. These data show that P10 HI increases P13 hippocampal Iba1 relative to Sham, and that there is a sex-dependent difference in this HI impact: Iba1 variance is greater in Male-HI vs Female-HI mice.

### 3.3. The CSF1R inhibitor PLX3397 depletes Iba1+ cells by P10 without disrupting body weight

Most prior work with PLX3397 has given the drug orally for two weeks or longer,^47,50^ a duration which is not practical for testing the role of microglia in P10 HI injury. Therefore, in Cohort 2 we tested if, compared to mice that received Veh injections, intraperitoneal injections of PLX3397 (either 25 or 50mg/kg) given daily P7-P9 would deplete brain Iba1+ cells by P10 **(Fig. 2A)**, the time when HI or Sham would be given to future cohorts. In regard to body weight, all three groups (Veh, 25mg/kg PLX, 50mg/kg PLX) were heavier on P10 than P7, showing they all gained weight over time. However, there were no weight differences among the groups at any time point (**Supp. Fig. 2**). In regard to Iba1+ cells, both mice receiving 25 and 50mg/kg PLX had fewer Iba1+ cells vs Veh mice. Qualitatively, even fewer Iba1+ cells were seen in 50mg/kg PLX vs 25mg/kg PLX mice **(Fig. 2)**. However, a pilot study showed the 50mg/kg dose of PLX led to unexpected mortality when combined with HI at P10 (data not shown; Cohort 3). Based on this mortality, all subsequent mice (Cohort 4) received 25mg/kg PLX or Veh. Quantitatively in Cohort 2, the density of Iba1+ cells was 97% lower in 25mg/kg PLX mice vs Veh mice **(Fig. 2)**. These data show that daily injections of 25mg/kg PLX from P7 to P9 robustly eliminate Iba1+ cells when examined at P10, the time point when we planned to perform HI.

**Figure 2.**
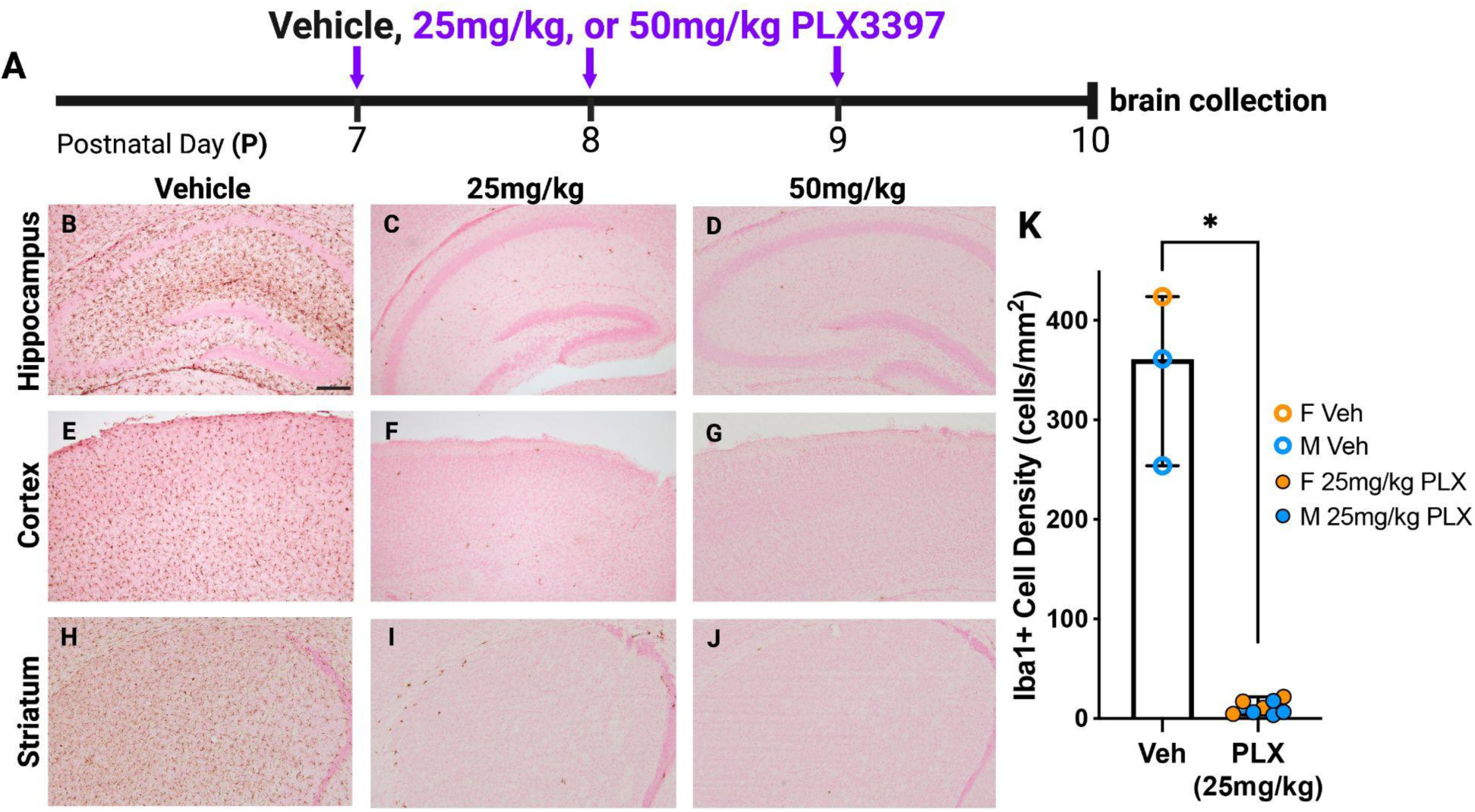
PLX3397 (25mg/kg, P7-P9) depletes Iba1-immunoreactive (+) cells when examined on P10. **(A)** <u>Timeline for Cohort 2:</u> Mouse pups received a daily *ip* injection of Veh, 25mg/kg PLX, or 50mg/kg PLX (indicated by purple arrows) from P7 to P9 and brains were collected 24 hours later on P10. **(B-J)** <u>Representative Iba1+ cell images</u> of coronal sections from the hippocampus, cortex, and striatum in Veh, 25mg/kg PLX, and 50mg/kg PLX mice. Magnification 100x; scale bar **(B)** 200 µm, applies to **(B-J)**. **(K)** <u>Iba1+ cell density</u> reflects the average of Iba1+ cell density in the striatum, cortex, and hippocampus of the left hemisphere in Veh and 25mg/kg PLX mice. Iba1+ cell density in 25mg/kg PLX mice was 97% less than Veh mice. There was more variance of Iba1+ cells in Veh vs 25mg/kg PLX mice. Complete details in **Table 1**.

### 3.4. Pup strength, exploration, and weight are not influenced by PLX, and HI does not influence exploration or weight

Having established that 25mg/kg PLX given P7 to P9 depletes microglia by P10, in Cohort 4 we next tested if microglial depletion through P13 alters fundamental behaviors, such as grip strength and exploration in a novel environment **(Supp. Fig. 3B-C)** and body weight **(Supp. Fig. 3D-E)**^53^. These mice also underwent HI or Sham at P10 **(Fig. 3)**. In the forelimb grasp test, a measure of strength, all groups grasped for longer at P9 than at P8 with no additional differences **(Supp. Fig. 3B)**. In open field traversal at P12, a test of exploratory behavior, all groups had similar latencies to exit the circle **(Supp. Fig. 3C)**. Weights were collected from P7 to P13, with negligible impact of PLX or HI on weight **(Supp. Fig. 3D)**. Analysis of only the final weight at P13 differed by Sex, but there was no *post hoc* difference **(Supp. Fig. 3E).** In regard to fur, two qualitative changes in PLX mice were noted **(Supp. Fig. 4)**. At P10 during surgery (not shown) and also at P13 **(Supp. Fig. 4A)**, PLX mice had a pinker abdomen vs Veh mice. At P13, PLX mice had notable hair loss on their snout, behind their ears, and along their fore- and hind paws vs Veh mice **(Supp. Fig. 4B)**, a pattern perhaps indicative of over-grooming. Aside from fur state and hair loss, at P13 PLX mice were qualitatively similar to Veh mice regarding movement and interactions with littermates in both the home cage and in a novel environment. These data show that PLX does not influence grasping, exploratory behavior, or weight, and that HI does not influence exploratory behavior or weight.

**Figure 3.**
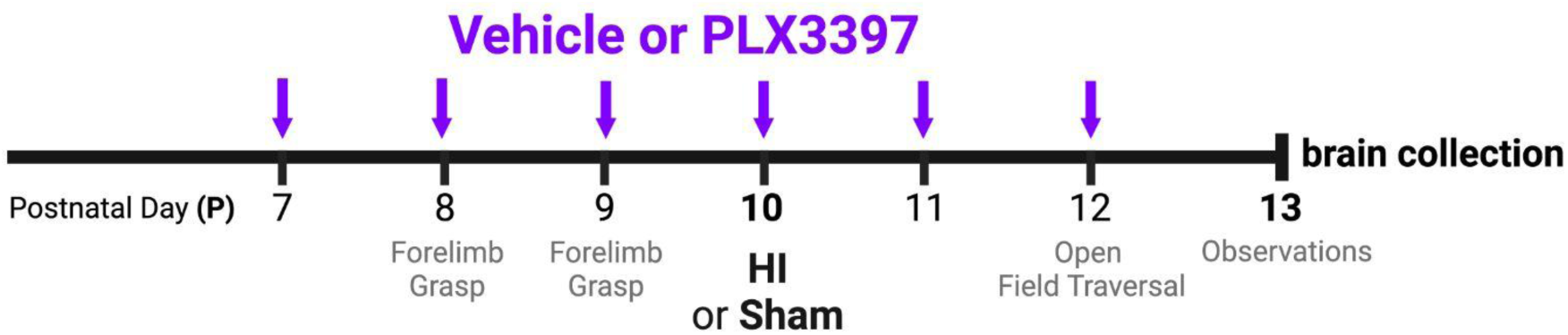
Timeline for Cohort 4 mice. Mice received daily intraperitoneal injections of Vehicle or PLX (25mg/kg, purple arrows) every 24 hours on P7-P12, and brains were collected 24 hours after the last injection on P13. At P10, mice underwent hypoxia-ischemia (HI) or Sham surgery and normoxia. Forelimb grasp (P8, P9) and open field traversal (P12) were assessed 30-45 min after the daily injection.

### 3.5. PLX does not influence HI-induced P10 seizure score

Although PLX did not influence behavior or weight between P7 and P13, it is reasonable to ask if PLX given P7-P10 prior to HI or Sham on P10 influenced an immediate effect of HI: seizure activity seen before, during, or after hypoxia. To this end, Cohort 4 P10 mice were observed after carotid ligation or Sham surgery and prior to, during, and for 10 min post-hypoxia for evidence of seizure activity. The average total seizure scores during hypoxia were similar among the four HI groups **(Supp. Fig. 5)**. Therefore, there was no effect of PLX (thus far given P7-P10) on HI-induced seizure activity.

### 3.6. In Female-HI mice, PLX increases the proportion of mice with pale brain surface lesion at P13, while in Male-HI mice, PLX decreases this proportion

PLX did not influence a short-term effect of HI (seizure activity on P10), yet it is possible that PLX given P7-P12 changed a longer-term effect of HI: the proportion of HI mice on P13 that have a pale lesion on the surface of the extracted brain. As expected, no Sham mice (either PLX or Veh) of either sex had a pale lesion on the brain surface, in keeping with our earlier results **(Supp. Fig. 1)**. Of the 51 HI mice that received PLX or Veh injections P7-P12, 12% (6/51) had a pale lesion on the surface of the brain at P13.

These six mice represented 8% of Veh-HI (2/25) and 15% of PLX-HI (4/26) mice, percentages which are not statistically different **(Table 1)**. When further broken down by sex, these six mice represented 0% of Female-Veh-HI (0/12), 23% of Female-PLX-HI (3/13), 15% of Male-Veh-HI (2/13), and 8% of Male-PLX-HI (1/13) mice. While these percentages did not statistically differ (Supp. Fig. 6, **Table 1)**, when visualized there was a notable sex-specific PLX effect **(Supp. Fig. 6)**. Specifically, in Female-HI mice, PLX increased the proportion of mice with pale brain surface lesion at P13, while in Male-HI mice, PLX decreased this proportion **(Supp. Fig. 6)**.

### 3.7. Iba1+ cell depletion worsens HI lesion scores in females but not males

As only a portion of HI mice of both sexes showed a pale lesion on the surface of the brain, we next utilized a more robust approach to assess HI-induced damage and the potential impact of sex and PLX: histopathologic lesion score of cresyl-violet stained brain sections.^60^ Observations of cortex and hippocampal subregions yielded lesion scores ranging from 0 (no injury) to 3 (severe cell loss with cystic infarction; **Fig. 4A-T**). Both hemispheres were examined. Stained sections from the contralateral (left) hemisphere of every mouse — regardless of sex, treatment, or surgery condition — yielded lesion scores of zero. In addition, the ipsilateral (right) hemispheres of all Sham mice yielded scores of zero. In the ipsilateral hippocampus of HI mice **(Fig. 4U)**, Female-PLX-HI mice had a higher average lesion score vs Female-Veh-HI mice and Male-PLX-HI mice. Male-PLX-HI mice had a visually lower average lesion score vs Male-Veh-HI mice, which was notably in the opposite direction from females; however, this visual trend was not statistically different **(Table 1)**. In the ipsilateral cortex **(Fig. 4V)**, there was no difference among the four groups. In addition to average lesion score, data were visualized by bins of lesion severity **(Fig. 4W, X)**. This alternate presentation supports the impression that PLX increases lesion scores in females and decreases lesion scores in males. Each ipsilateral hippocampal subregion was consistent **(Fig. 4Y-BB)**: lesion score analysis suggesting that PLX had an opposite effect in Female-HI and Male-HI mice. For example, in CA1 **(Fig. 4Y)**, Male-PLX-HI mice had a lower average lesion score than Male-Veh-HI mice and Female-PLX-HI mice. In CA2 **(Fig. 4Z)**, Female-PLX-HI mice had a higher average lesion score vs Female-Veh-HI mice and Male-PLX-HI mice. In hippocampal CA3 **(Fig. 4AA)**, Female-PLX-HI mice had a higher average lesion score vs Female-Veh-HI and Male-PLX-HI mice. The only sampled brain regions where PLX did not influence lesion size were the hippocampal DG **(Fig. 4BB)** and the striatum (data not shown). After determining lesion scores for each brain region, each HI mouse was then assigned an average ipsilateral lesion score, and this was compared to the seizure score (from P10) to assess any relationship. There was no correlation between the P10 seizure score and average ipsilateral lesion score **(Supp. Fig. 7)**. In sum, these data suggest that the effects of PLX-mediated Iba1+ cell depletion on HI-induced lesions are opposite in females and males: PLX worsens lesions in Female-HI mice, but ameliorates lesions in Male-HI mice.

**Figure 4.**
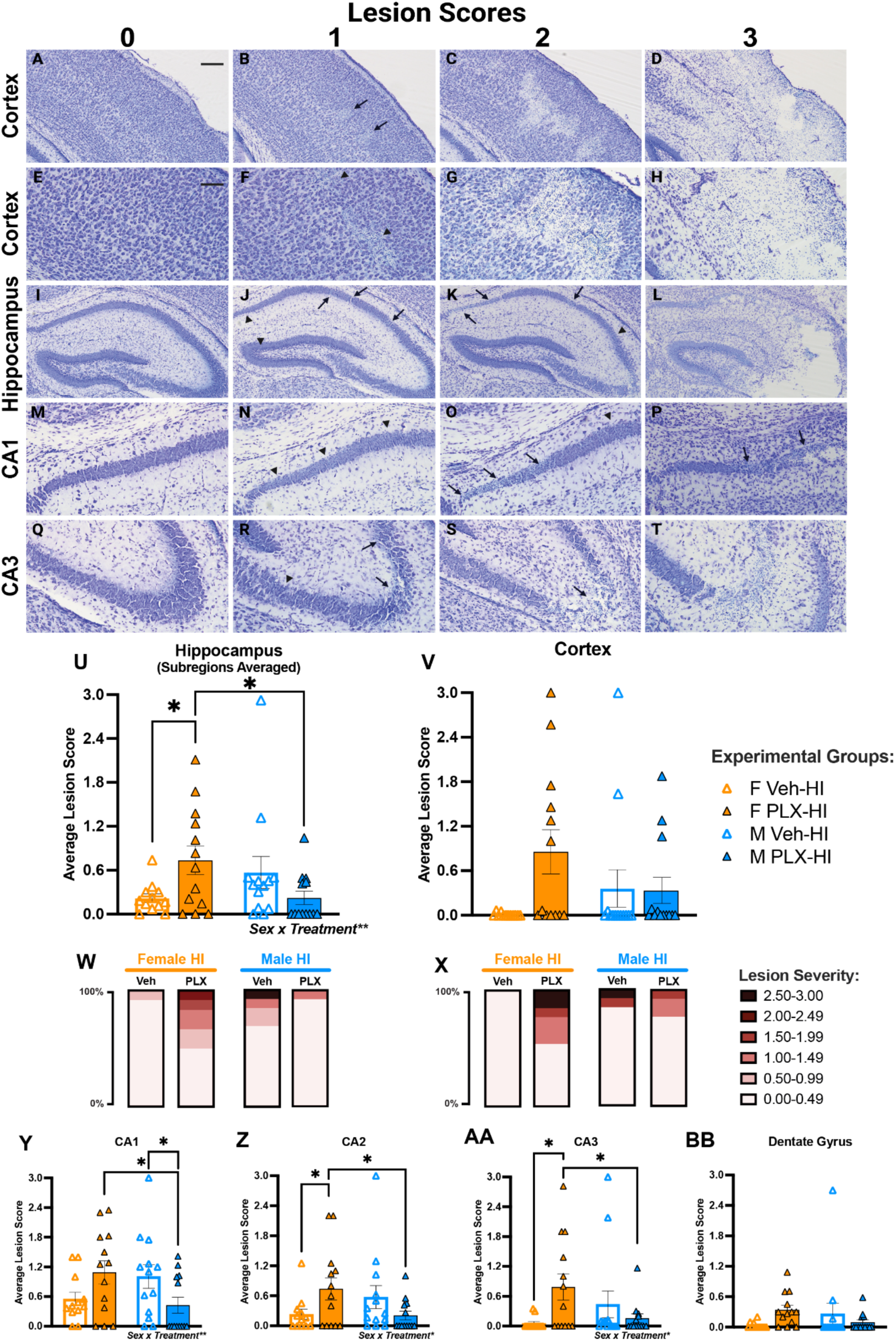
PLX worsens lesion score in HI female mice but not in HI male mice. **(A-T)** <u>Qualitative assessment</u> of HI-induced lesions via cresyl violet-stained cortical and hippocampal sections from Cohort 4 mice. 0=no injury, 1=scattered shrunken neurons, 2=moderate cell loss, 3=severe cell loss with cystic infarction. Arrowheads=shrunken neurons; arrows=cell loss. Scale bar **(A)**=200 µm, applies **(A-D)**,**(I-L); (E)**=100 µm, applies **(E-H), (M-T)**. **(U-BB)** <u>Average lesion score</u> for the **(U)** hippocampus, **(V)** cortex, **(Y)** CA1, **(Z)** CA2, **(AA)** CA3, and **(BB)** dentate gyrus and <u>binned lesion severity</u> for the **(W)** hippocampus and **(X)** cortex in all Cohort 4 mice. Female-PLX-HI mice had higher average lesion scores than Female–Veh-HI and Male-PLX-HI in the hippocampus, CA2, and CA3 **(U,Z,AA)**. Male-PLX-HI had lower average lesion scores than Female-PLX-HI and Male-Veh-HI in CA1 **(Y)**. F, Female. HI, hypoxia ischemia. M, Male. PLX, PLX3397. Veh, Vehicle. Complete details in **Table 1**.

### 3.8. PLX depletes Iba1+ cells in female and male mice equally

One possible explanation for these disparate effects — PLX makes the lesion worse in Female-HI mice but has the opposite effect in Male-HI mice — is that PLX depletes Iba1+ cells unequally in female vs male mice. To test this in our large experimental cohort, Iba1+ cells in 3 brain sections from every mouse in Cohort 4 including both PLX and Veh mice were qualitatively and quantitatively assessed.

Qualitatively, the depletion after PLX treatment was dramatic and readily apparent throughout all brain regions analyzed, including cortex, hippocampus, striatum, and thalamus. While in Veh mice Iba1+ cells were evenly tiled densely throughout the brain, in PLX mice there were far fewer Iba1+ cells and their distribution was irregular. An observer masked to the injection group could easily distinguish Iba1-stained sections between Veh **(Fig. 5A)** vs PLX mice **(Fig. 5B)**. There was also a qualitative morphological difference in the few residual Iba1+ cells in PLX mice vs Iba1+ cells in Veh mice. For example, the residual Iba1+ cells in PLX mice were larger and thicker than typical Iba1+ cells seen in Veh mice, and they predominated in white matter tracts rather than gray matter areas.

**Figure 5.**
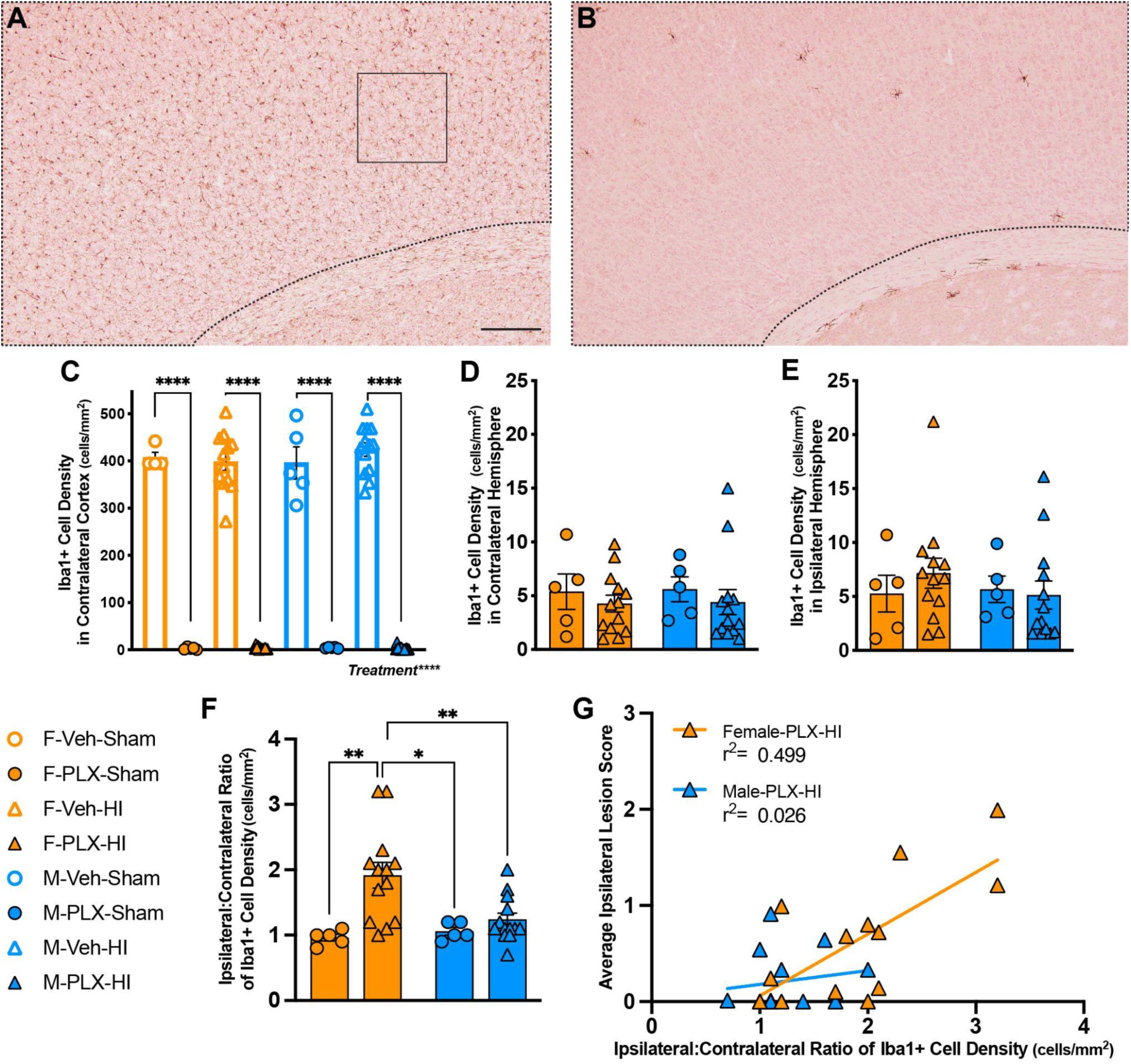
PLX depletion of Iba1+ cells is not sex-dependent. **(A-B)** <u>Representative</u> <u>photomicrographs</u> of Iba1-stained cortical sections from Female-Sham mice that received Veh **(A)** or PLX (**B**; 25mg/kg *ip*, daily P7-P12). Scale bar **(A)**=200 µm, applies to **(A-B)**. Note lack of brown Iba1+ cells after PLX **(B)** vs Veh **(A)**. Dotted lines **(A,B)**: cortical area analyzed for Iba1+ cell density. Solid outlined square in **(A)**: 400×400 pixel unit where Iba1+ cells were manually counted in Veh mice. **(C)** In Cohort 4 mice, PLX mice had 115-fold lower <u>density of Iba1+ cells in the contralateral cortex</u> than Veh mice, equivalent to a >99% depletion of Iba1+ cells. **(D, E)** In Cohort 4 PLX mice, there was no difference in <u>Iba1+ cell density</u> among the four PLX groups in either the <u>contralateral</u> <u>hemisphere</u> **(D)** or the <u>ipsilateral hemisphere</u> **(E). (F)** Female-PLX-HI had larger <u>ipsilateral:contralateral hemisphere ratio of Iba1+ cell density</u> than Female-PLX-Veh, Male-PLX-HI, and Male-PLX-Veh. **(G)** <u>Ipsilateral:contralateral ratio of Iba1+ cell density</u> correlated with the <u>average ipsilateral lesion score</u> in Female-PLX-HI mice but not in Male-PLX-HI mice. F, Female. HI, hypoxia ischemia. M, Male. PLX, PLX3397. Veh, Vehicle. Complete details in **Table 1**.

Quantitatively, PLX robustly decreased Iba1+ cells: PLX mice had 115-fold lower density of cortical Iba1+ cells than Veh mice **(Fig. 5C)**, a >99% depletion of Iba1+ cells. There was no effect of Sex or HI on Iba1+ cell density **(Fig. 5C)**. To assess for a potential sex difference in PLX-induced Iba1+ cell depletion, we manually counted in all PLX mice the few Iba1+ cells remaining in each hemisphere across 3 different sections. The Iba1+ cell density was similar across both sexes and HI or Sham condition in the contralateral hemisphere **(Fig. 5D)** and the ipsilateral hemisphere **(Fig. 5E)** of PLX mice.

However, in the PLX mice, the ratio of Iba1+ cell density in the ipsilateral:contralateral hemispheres was higher in the Female-HI mice than the Female-Sham, Male-Sham and Male-HI mice **(Fig. 5F)**. The average ipsilateral lesion score did not correlate with the Iba1+ cell density in either the contralateral or ipsilateral hemisphere in Female-PLX-HI and Male-PLX-HI mice **(Table 1)**. However, the average ipsilateral lesion score did correlate with the ratio of the Iba1+ cell density in the ipsilateral:contralateral hemispheres in the Female-PLX-HI but not the Male-PLX-HI mice **(Fig. 5G)**. Thus, with r^2^=0.499 in Female-PLX-HI mice, half of the variance in the ipsilateral lesion score of this group is explained by the higher Iba1+ cell density in the ipsilateral vs contralateral hemisphere. Qualitatively, we did not see Iba1+ cells massively congregating around lesion sites in any of the PLX-HI mice; this is in contrast to Veh-HI mice that have obvious perilesional clusters of Iba1+ cells.

## 4. DISCUSSION

We report two major findings. First, robust pharmacologic depletion of brain microglia before P10 is possible in mice using daily *ip* injections of PLX3397 (P7-P9). Compared with giving this class of drugs orally over weeks, this method yields effective, rapid depletion that allows greater temporal precision to study the impact of microglia depletion in young animals. Second, PLX-induced depletion of Iba1+ cells prior to P10 HI leads to a worse HI lesion score in Female mice vs Veh Female mice, but not in Male mice; in fact, Male HI lesion scores appear to be improved by PLX vs Veh. This sex-dependent effect of PLX on HI-induced lesion score is not due to a sex-dependent effect of PLX on global Iba1+ cell depletion.

The immune response to HI involves cells that originate from two compartments: centrally-originating cells primarily consisting of brain-resident microglia and peripherally-originating cells including monocyte-derived macrophages (MDM cells) that infiltrate the brain after injury. The uninjured brain is thought to contain few MDM cells, so Iba1+ cells in the brain parenchyma of uninjured mice are generally considered to be microglia^63^. We show in females that HI lesions are made worse when microglia are depleted, indicating that microglia are protective in female brains. In contrast, microglial depletion in males tends to make HI lesions smaller. There are several possible explanations for these sex-specific effects, two of which we will expand on here. First, it is possible that in females microglia are protective and in males microglia are not protective. Alternatively, it is possible that in females protective microglia are the main macrophage response to HI, but that in males the peripheral MDM response to HI overpowers the protective capacity of central microglia. Supporting this latter speculation, males after HI have more MDM cells infiltrating the brain than females.^5^ We show that both males and females have more Iba1+ cells 3 days after HI vs Sham, but that males have more variability in the magnitude of their response (**Fig. 1**). We hypothesize that the brain immune response to HI has underlying latent sex differences — when a similar-appearing outcome is achieved through distinct mechanisms.^64^ Specifically, while sex differences in Iba1+ cell number are subtle after HI in microglia-intact brains, depleting microglia prior to HI reveals the differing roles these Iba1+ cells have in male and female brains.

Our results are very interesting in the context of the literature. Two prior studies tested the effect of microglial depletion on HI-induced lesions and came to opposite conclusions. One study, like us, used a pharmacological approach (PLX3397, 25mg/kg orally) to deplete microglia and found smaller HI-induced lesions in microglia-depleted mice.^46^ However, they administered PLX3397 orally for 8 days (P4-P11, HI at P9), in contrast to our shorter 5-day *ip* administration of PLX (P7-P12, HI at P10). Also notable given our sex-specific effects of microglia depletion on HI lesion is that this study did not disaggregate their data by sex. The other study did disaggregate their data by sex, and found that microglial depletion via inducible genetic modification (Cx3cr1^CreER/+^ Rosa26^DTA/+^ mouse) increased the size of HI lesions in male — but not female — mice.^42^ This is in contrast with our data, which show microglial depletion increased the HI-induced lesion in female — but not male — mice.

There are at least two possibilities as to why our work shows microglia are protective in female (but not male) mice, while the inducible genetic depletion approach shows microglia are protective in male (but not female) mice.^42^ First, as the authors themselves note, the use of tamoxifen in their inducible genetic model may lead to a neuroprotective effect of this estrogen-like compound predominantly in female mice.^42^ A second hypothesis for our conflicting results is that their depletion approach may amplify the peripheral immune response, which — as speculated above — may contribute to HI lesion in males more than females. The two different approaches to deplete microglia (pharmacologic vs inducible genetic) have different effects on peripheral monocytes and this likely contributes to the opposite results. Specifically, the inducible genetic approach increases the number of Ly6C^hi^ monocytes in the spleen, but the pharmacologic approach we utilize, PLX3397, does not change the number of these cells.^47,65^ It is possible the inducible genetic approach may amplify the peripheral inflammatory component of HI, yielding larger injury when peripheral MDM cells are called into the brain, and thus affecting males more than females. In contrast, if the female Iba1+ cells after HI are predominantly central microglia, then the female response to injury may be less impacted by worsened peripheral inflammation. This hypothesis is highly testable in future studies.

Aside from the role of peripheral cells, our results here clearly demonstrate that brain resident microglia have a protective role in female mouse brains. In addition to peripheral monocytes, both genetic and pharmacologic depletion strategies have varying effects on other immune cells including lymphocytes, whose role is important to investigate^66^. PLX3397 may also impact oligodendrocyte precursor cells^67^. Route of administration (intraperitoneal, oral, intracisternal), dose, and timing all impact PLX’s effects on non-microglial cells.^47^ Studies specifically detailing sex differences in the relative contribution of microglia vs MDM cells after HI will be useful, as well as evaluating sex differences in the intrinsic properties of microglia, and in how MDM cells are called into the brain after injury. In general, these discrepant results highlight the importance of including both male and female mice in every study and analyzing data separately in each sex, as the experimental manipulations may have opposite effects in each sex that could negate each other and create an illusion of no effect. Latent sex differences may not be revealed by HI alone, but may come to light after microglial depletion combined with HI.

Taken together, our findings are consistent with a hypothesis that after HI, brain-resident female microglia act in a protective manner to mitigate brain injury, while the Iba1+ cells in male brains may consist of more MDM cells infiltrating from the blood and worsening the injury. If this is true, then blocking MDM cells from entering the brain may prove to be an effective therapeutic target. This perspective warrants further investigation.

## 5. CONCLUSIONS

In this study, we established a model of pharmacologic depletion of Iba1+ cells in young mice. We find that depletion worsens HI-induced lesions in females but not in males. Our findings indicate that Iba1+ cells play a neuroprotective role in female mice after hypoxia-ischemia, but that this may be different in male mice.

## List of abbreviations

Contra: contralateral
CV: cresyl violet
DG: dentate gyrus
F: Female
g: grams
HI: Hypoxia Ischemia
HIE: Hypoxic Ischemic Encephalopathy
IHC: immunohistochemistry
Ipsi: ipsilateral
M: Male
MDM: monocyte-derived macrophage
P: Postnatal Day
PLX: 25mg/kg PLX3397
RT: room temperature
s: seconds
v: volume
Veh: Vehicle

## DECLARATIONS

### Ethics approval and consent to participate

All mice were cared for in compliance with Protocol #1245 approved by the Institutional Animal Care and Use Committee at the Children’s Hospital of Philadelphia and guidelines established by the NIH’s *Guide for the Care and Use of Laboratory Animals*.

### Consent for publication

Not applicable

### Availability of data and materials

All data generated or analyzed during this study are included in this published article and its supplementary information files. The data are available from the corresponding author, DGB, upon reasonable request.

### Competing interests

The authors declare that they have no competing interests.

### Funding

This work was supported by the National Institutes of Health (CNCDP-K12 NS098482, R01MH129970, R01DK135871), CHOP Research Institute, CHOP Division of Neurology and philanthropic funds. Funding for biostatistical support provided by NICHD (P50HD105354) for the CHOP/Penn Intellectual and Developmental Disabilities Research Center.

### Authors’ contributions

Conceptualization: DGB

Data curation: DGB, SNG

Formal Analysis: DGB, SNG

Funding acquisition: DGB, AJE

Investigation: DGB, SNG, KJJ

Methodology: DGB, SNG, KJJ

Project administration: DGB

Resources: DGB, SY, AJE

Supervision: DGB, SY, AJE

Validation: DGB, SNG

Visualization: DGB, SNG, SY, AJE

Writing, original draft: DGB, SNG

Writing, review & editing: DGB, SY, AJE, SNG

## Acknowledgements

We would like to thank: Oscar Marcos-Contreras (UPenn) and Raul Chavez-Valdez for teaching DGB how to perform HI; Sheyenne Gillis for assistance setting up the HI model and collecting data for first cohort of HI mice; Ida Narli for assistance with Iba1 signal quantification in PLX-HI cohort; F. Chris Bennett, Sonia Lombroso, and Mariko Bennett for helpful discussions about microglia depletion; Fred Kiffer, Lorianna Colon and Ruthie Wittenberg for suggestions on improving the manuscript and clarity of figures; Fred Kiffer for 3D printing equipment for developmental milestones; Rui Xiao, Antoneta Karaj and Gabrielle Gionet for statistical guidance.

## SUPPLEMENTARY FIGURES

**Supplementary Figure 1.**
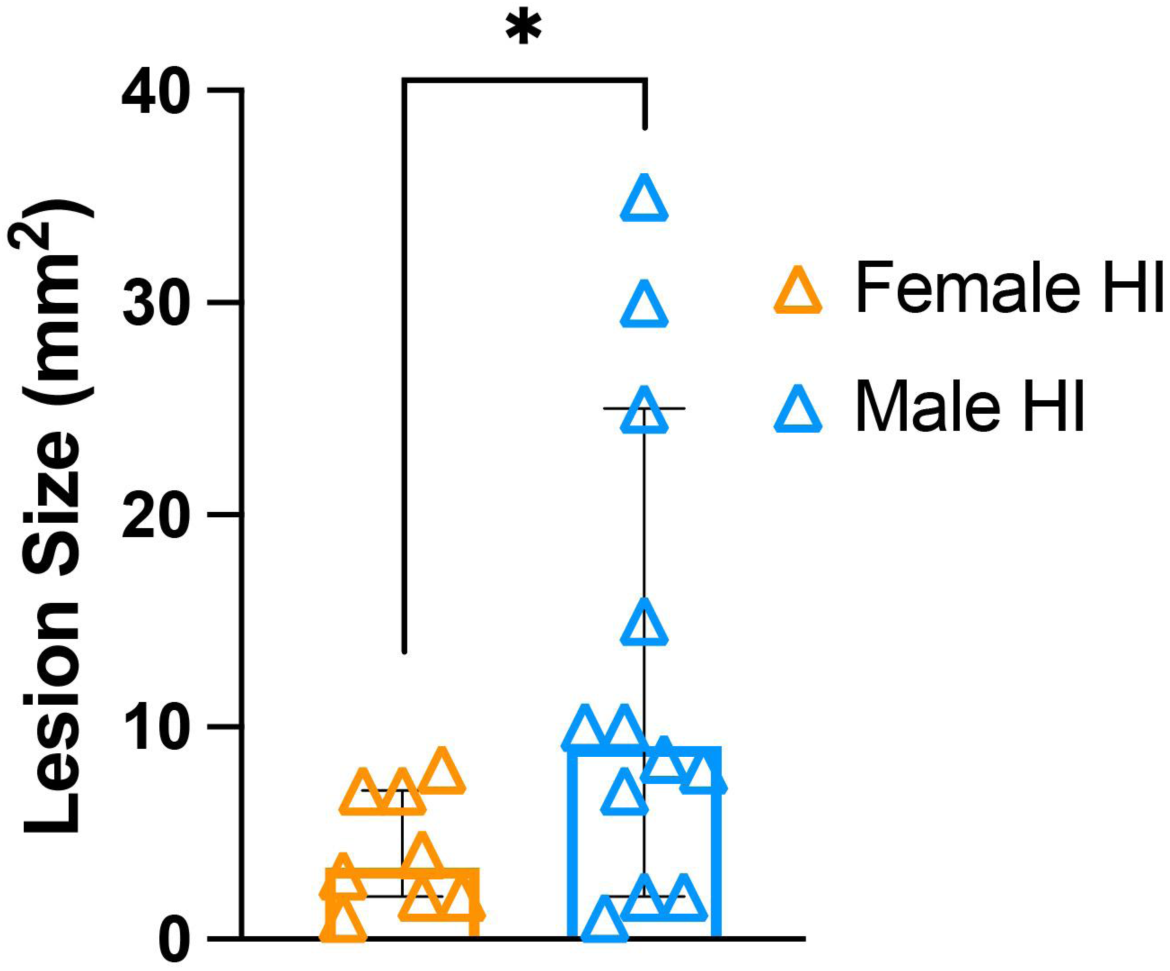
At P13, a portion of HI mice had a pale lesion on the brain surface, and the lesion was larger in Male-HI vs Female-HI mice. <u>Size of pale</u> <u>lesion on the brain surface</u> in whole brains from P13 Male-HI and Female-HI mice at P13 (Cohort 1). No lesions were seen in Sham mice or in the majority of HI mice, and those zero values are not shown here. Pale lesions on the brain surface of Male-HI mice were larger vs Female-HI mice. Additionally, there was more variance of lesion size in Male-HI than Female-HI mice. Complete details in Supplementary Table 1 (Supp. Table 1).

**Supplementary Figure 2.**
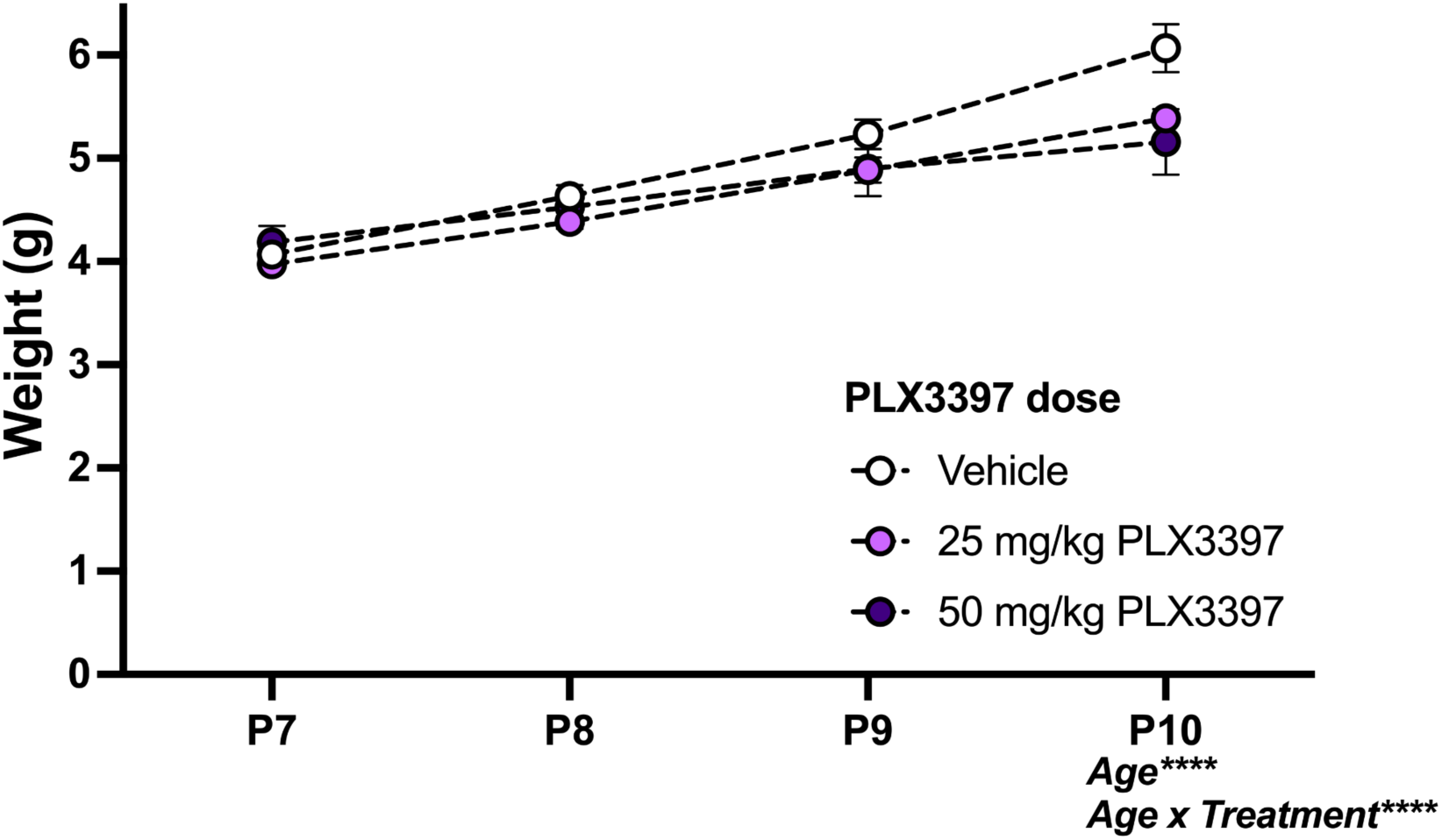
Daily injections of PLX3397 from P7 to P9 did not affect weight gain. <u>Weights of Cohort 2 mice from P7 to P10</u>. Each group was heavier on P10 vs P7, showing mice in each group gained weight over time. Despite visual difference, there was no difference in the weights among the three groups at any time point. Complete details in **Supp. Table 1**.

**Supplementary Figure 3.**
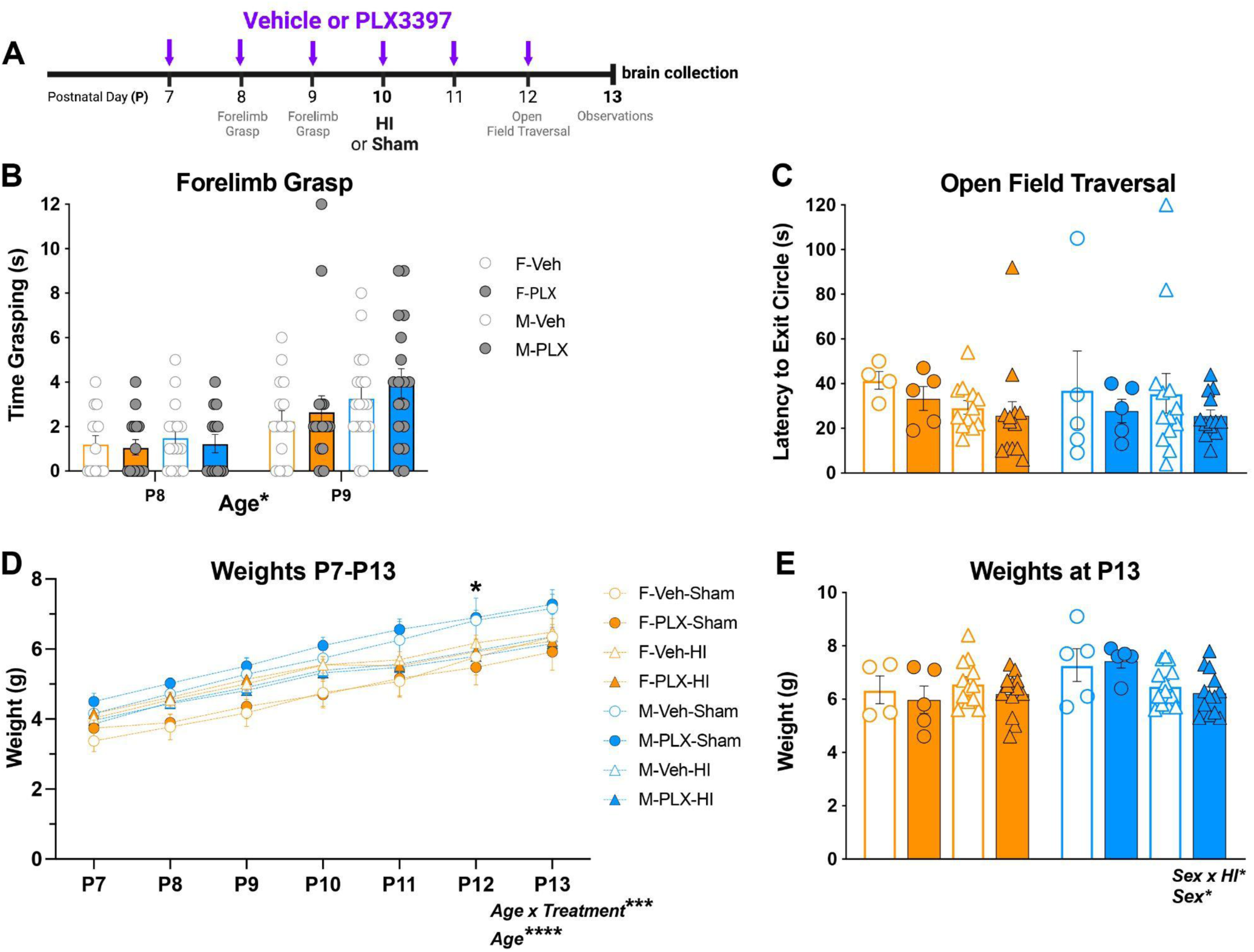
Neither PLX treatment (P7-P12) nor HI (P10) affects strength, exploration, or weight at P13. Negligible impact of PLX or HI on weight from P7 to P13. **(A)** Timeline for Cohort 4 mice. Forelimb grasp and open field traversal were assessed 30-45 min after the daily ip injection (purple arrows) of PLX (25mg/kg) or Veh. **(B)** <u>Forelimb grasp</u> was performed on P8 and P9, prior to HI or Sham surgery. Therefore, mice are grouped by sex (Female: orange outline/fill; Male: blue outline/fill) and injection (Veh: gray-open circle; PLX: gray-filled circle). P9 mice grasped longer than P8 mice. **(C)** <u>Open field traversal</u> was performed on P12. All groups had similar times in P12 open-field traversal (latency to exit circle). **(D-E)** <u>Body weights</u> assessed daily P7 through P13 **(D)** showed all groups gaining weight. At P12, Male-PLX-HI weighed 16% less than Male-PLX-Sham. **(E)** At P13, there was a main effect of Sex and an interaction of Sex and HI on body weight. F, Female. g, grams. HI, hypoxia ischemia. M, Male. PLX, PLX3397.s, seconds. Complete details in **Supp. Table 1**.

**Supplementary Figure 4.**
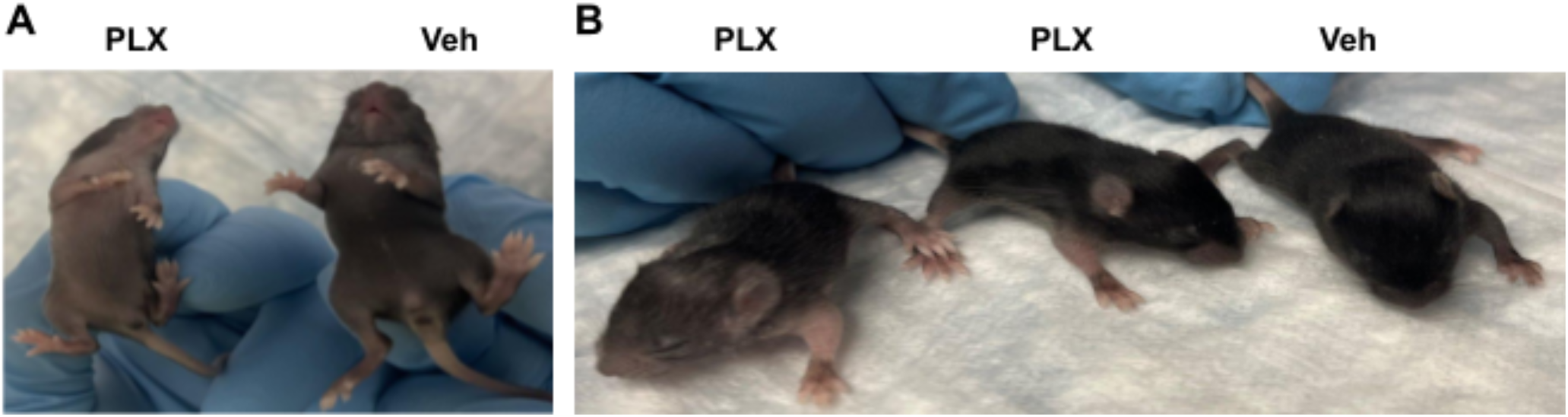
Qualitative analysis at P13 shows PLX mice have differences in fur state and hair loss compared to Veh mice. **(A)** After PLX or Veh injection daily P7-P12, P13 PLX mice had a pinker abdomen than Veh mice, which was also seen on the day of surgery (P10, images not shown). **(B)** P13 PLX mice also had less hair on their snout, behind their ears, and along their fore- and hind paws compared to Veh littermates. Left-most mouse in **(B)** is one of the most severe cases of dorsal surface hair loss. There was no qualitative difference between PLX-HI and PLX-Sham mice of either sex in either the color of the abdomen or the amount of hair loss on the dorsal surfaces of the body. PLX, PLX3397. Veh, Vehicle.

**Supplementary Figure 5.**
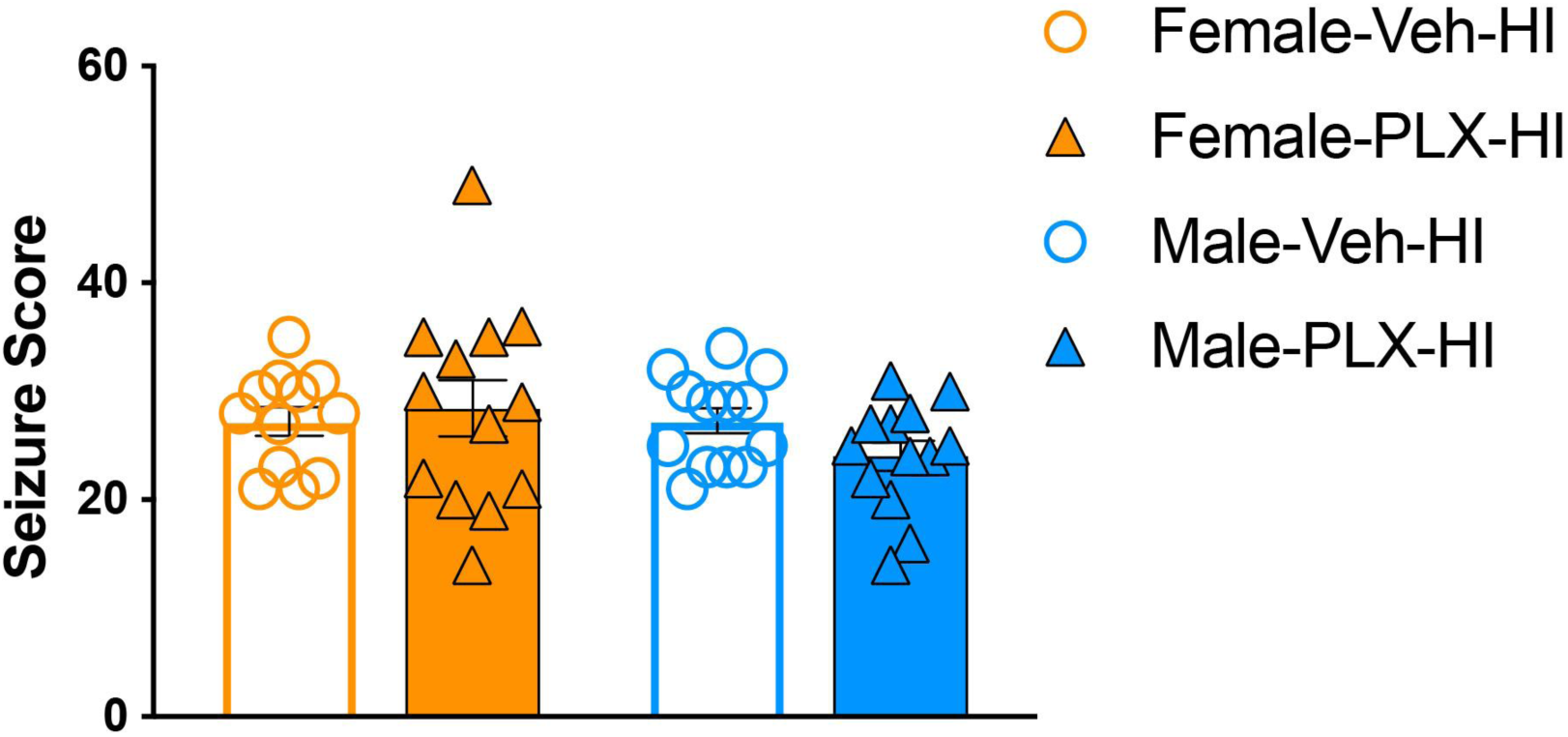
PLX does not impact P10 seizure score. <u>Seizure scores</u> from Veh or PLX (daily injections, P7-P10) Female or Male mice on day of Sham or HI (P10). Seizure scores are shown for HI mice during hypoxia; data for Sham mice are not shown as they all had seizure scores of zero. The average total seizure scores during hypoxia were not different among the four groups. F, Female. HI, hypoxia ischemia. M, Male. PLX, PLX3397. Veh, Vehicle. Complete details in **Supp. Table 1**.

**Supplementary Figure 6.**
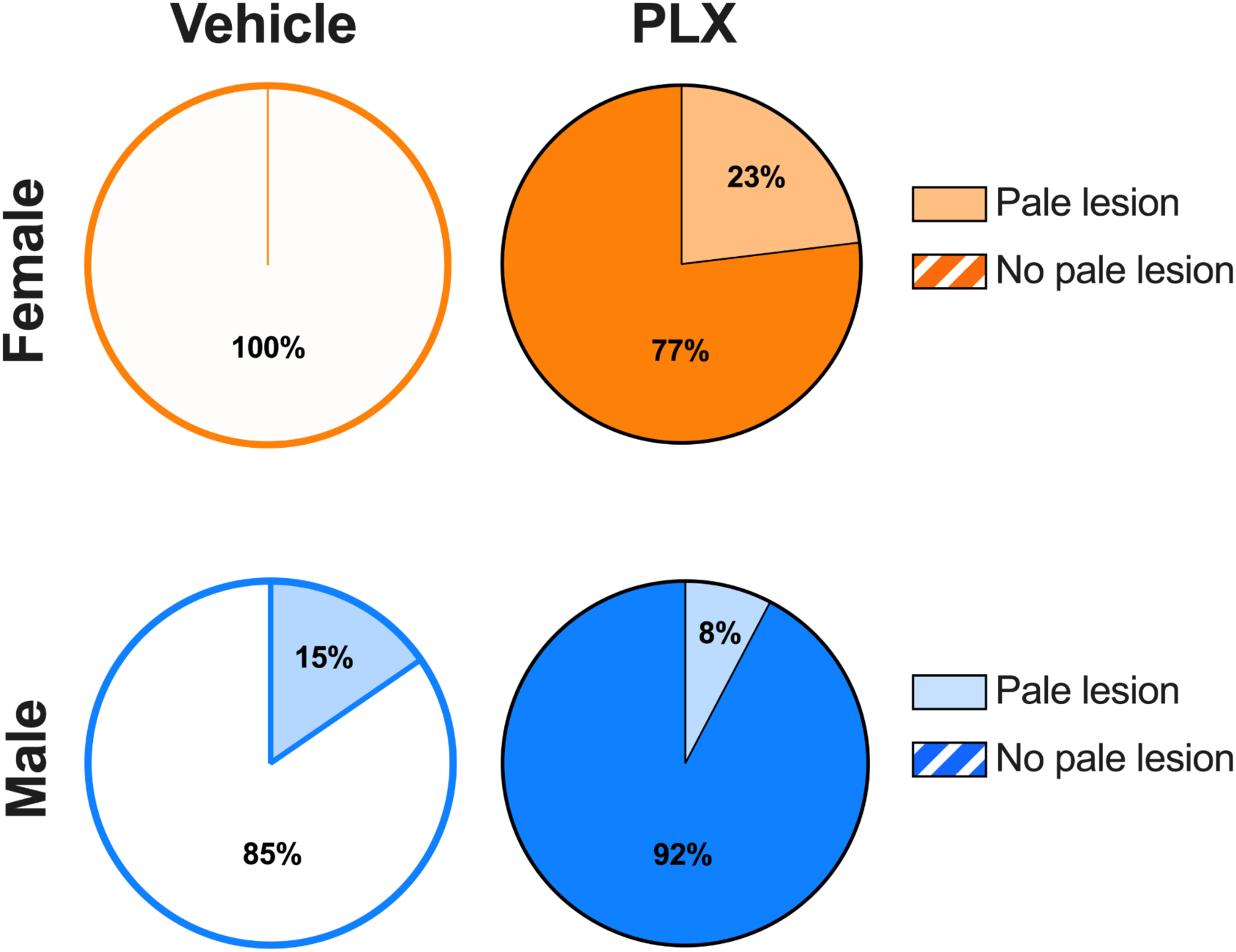
Among Veh or PLX mice who underwent HI, the fraction of mice per group with a pale lesion visible on the surface of the brain at P13 is differentially influenced by sex. <u>Fraction of mice with a pale lesion visible on the brain</u> <u>surface at P13.</u> Of the 52 mice who received either PLX or Veh and HI, 6 mice had a pale lesion on the surface of the brain. These six mice consisted of: 0% of the Female-Veh-HI mice (0/12), 23% of Female-PLX-HI mice (3/13), 15% of Male-Veh-HI mice (2/13), and 8% of the Male-PLX-HI mice (1/13). There was no association between Treatment and fraction of mice with pale lesions. PLX, PLX3397. Veh, Vehicle. Complete details in **Supp. Table 1**.

**Supplementary Figure 7.**
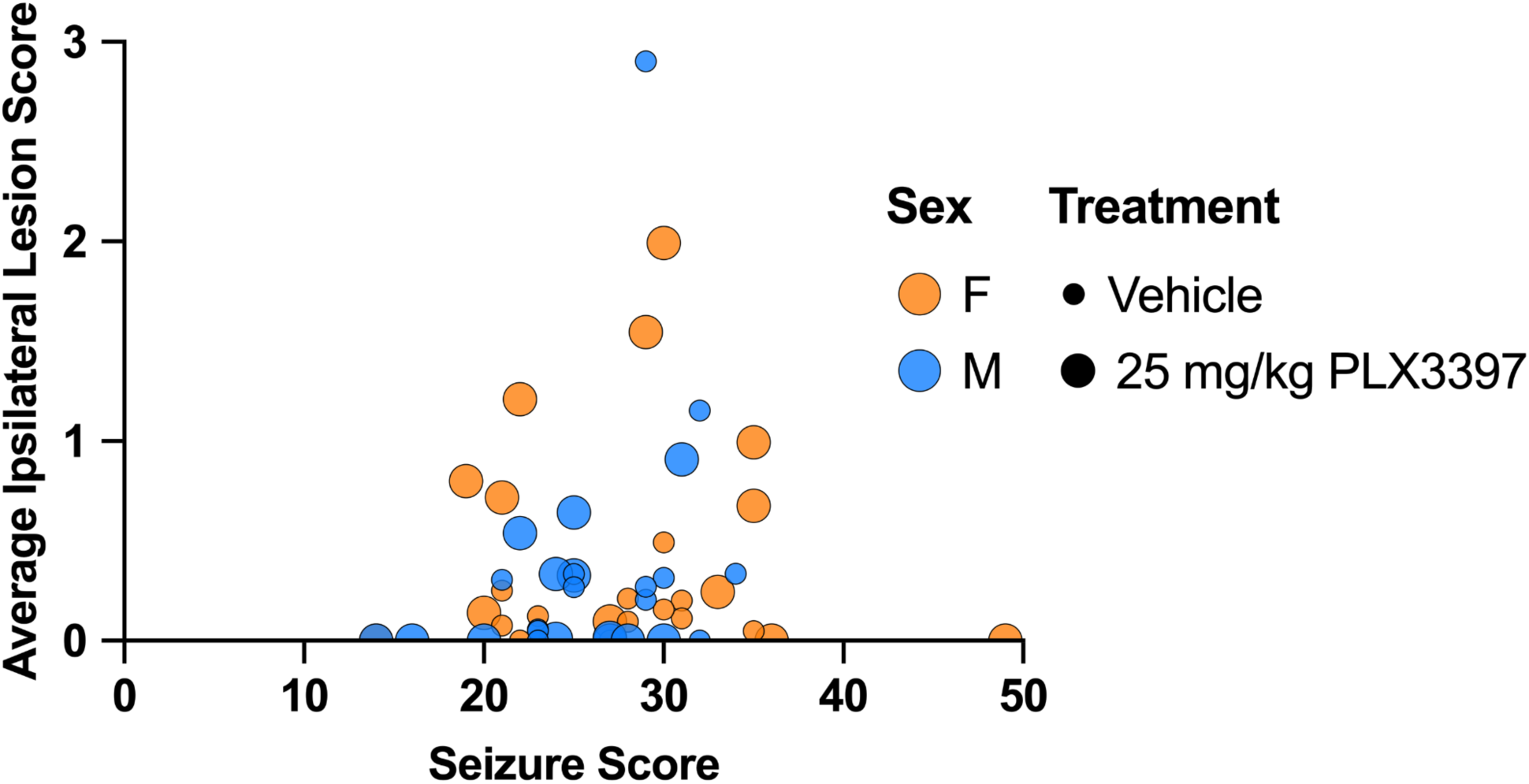
In HI mice, average ipsilateral lesion score does not correlate with seizure scores. Mice in Cohort 4 received daily PLX or Veh injections from P7 to P12 and HI or Sham on P10. Brains were collected on P13. Sham mice are omitted from this graph since they all had seizure scores of zero and lesion scores of zero. <u>Seizure scores (at P10) and average ipsilateral lesion score (determined by</u> <u>analysis of cresyl violet staining from brains collected on P13)</u> are graphed here. There was no relationship between average ipsilateral lesion score and seizure score when comparing Sex (Female, orange; Male, blue) and Treatment (Veh, smaller icon; PLX, larger icon) F, Female. M, Male. Complete details in **Supp. Table 1**.

## SUPPLEMENTARY MATERIALS AND METHODS

### S.2.1. Animals

Dams and males were C57BL/6J mice shipped from Jackson Laboratory and bred in the Children’s Hospital of Philadelphia (CHOP) Research Institute. Dams were individually housed in a HEPA-filtered, closed airflow vivarium rack (Lab Products Inc., Enviro-Gard™ III) under a normal 12:12-hour light-dark cycle (06:00 on, 18:00 off) with *ad libitum* access to food (standard rodent chow, Lab Diets 5015 #0001328) and water. The AAALAC-accredited facility at the CHOP Research Institute is temperature-(22°C) and humidity-controlled (30-70%). Dams were housed with males for an average of 11 days before being singly-housed when pregnant. Cages were checked for pups once daily in the late afternoon. Mouse pups were kept in the same cage as the dam and their littermates following birth and were not disturbed for 6 days. All mice were cared for in compliance with Protocol #1245 approved by the Institutional Animal Care and Use Committee at CHOP and guidelines established by the NIH’s *Guide for the Care and Use of Laboratory Animals.* Our scientific reporting adheres to the ARRIVE 2.0 guidelines.^52^

### S.2.2. PLX3397 injections

PLX3397 (PLX, Selleck, Catalog No.S7818) was resuspended in dimethyl sulfoxide (Cell Signaling Technology, Catalog No.12611P) to make a 50mM stock, which was frozen and stored (-80°C) in aliquots of 50μL and 100μL. At P7, male and female littermates were randomly divided into groups. Mice received daily ∼9AM intraperitoneal injections (*ip*) of 20μL PLX or injections of the same volume 20μL Vehicle (50% phosphate-buffered saline PBS, 50% polyethylene glycol). Injection quality was noted each day for each mouse, and most days in most mice there was no residue on abdomen after *ip* injection.

### S.2.3. Experimental groups

Four cohorts of mice were included in this study. In each cohort, mice of each sex were randomly assigned to experimental groups evenly distributed within each litter. <u>Cohort 1</u> received no drug treatment and only underwent HI or Sham at P10 for assessment of Iba1 response at P13; this included 18 litters with 140 pups total: 63 females (20 Sham, 43 HI) and 77 males (25 Sham, 52 HI). Cohort 1 mortality rate was 4% (n=6): n=4 Male-HI and n=2 Female-HI; all of these pups died on P10. Cohort 1 pups were weighed at P10 and P13. A subset of these Cohort 1 mice were used for immunohistochemistry: Female-Sham n=6, Female-HI n=15, Male-Sham n=9, Male-HI n=23.

<u>Cohort 2</u> was used for a PLX3397 dose-response assessment with pups randomly assigned to three experimental groups: 50mg/kg PLX, 25mg/kg PLX, and Veh. Mice received daily ip injections from P7 to P9. Brains were collected at P10, 24 hours after the last PLX or Veh injection. Cohort 2 included 2 litters with 18 pups total: 3 Veh (1 female, 2 male), 8 mice receiving 25mg/kg PLX3397 (4 female, 4 male), and 7 mice receiving 50mg/kg PLX3397 (3 female, 4 male). One additional mouse in this cohort was an outlier due to low weight (21% lower than mean for her litter) so was excluded from all analysis. Cohort 2 mortality was zero. Cohort 2 pups were weighed daily from P7 to P10.

<u>Cohort 3</u> included one litter of 9 mice, 5 of which received 50mg/kg PLX3397 (2 Female-HI, 1 Female-Sham, 2 Male-HI) and 4 Veh mice (1 Female-HI, 1 Female-Sham, 2 Male-HI). Of these mice receiving 50mg/kg PLX mice, 80% died by P12 (100% of females including 2 HI and 1 Sham; 50% of HI males); there were no deaths in the Veh groups. Cohort 3 pups were weighed daily from P7 to P13 or until death.

<u>Cohort 4</u> received either 25mg/kg PLX3397 or Veh given *ip* P7-P12; brains were collected on P13, 24 hours after their last 25mg/kg PLX or Veh injection. This included 10 litters with 78 pups total: 38 females, 40 males; 10 Veh-Sham, 28 Veh-HI, 10 PLX-Sham, 30 PLX-HI. Cohort 4 had a 9% mortality rate (n=7): 14% (n=1) of deaths were Female-Veh-Sham, 14% (n=1) of deaths were Female-Veh-HI, 14% (n=1) of deaths were Female-PLX-HI, 29% (n=2) of deaths were Male-Veh-HI, and 29% (n=2) of deaths were Male-PLX-HI. There were no deaths in the Female-PLX-Sham, Male-Veh-Sham, or Male-PLX-Sham groups. Cohort 4 pups were weighed daily from P7 to P13.

### S.2.4. Behavioral Testing

Developmental milestones and behavior were tested on postnatal days 8, 9, 12, and 13 as previously published.^53^ At P8 and P9, forelimb grasp (strength) was measured using a custom-made 3D-printed forelimb grasp apparatus.^68^ Pups were placed vertically 10cm over the table surface and forepaws were allowed to grip a 0.80 mm diameter rod (straightened paper clip); the length of time the pup gripped the rod was recorded.^53^ At P12, exploration was assessed by placing each mouse in the center of a 13cm diameter circle (drawn on white paper) and recording the time until the pup exited the circle (latency to exit circle) or 120 seconds (s) elapsed.^53^ Both forelimb grasp and exploration were assessed 30-45 minutes (min) after the daily *ip* injection (purple arrows) of PLX or Veh.

### S.2.6. P10 Hypoxia-Ischemia

A modified Vannucci Model^2,54–56^ of HI was used in male and female PLX and Veh mice at P10. Male and female pups were randomly assigned to HI or Sham groups, ensuring as equal distribution as possible among all experimental groups within each litter.

Briefly, lidocaine ointment was placed on the pup’s ventral neck, then under isoflurane anesthesia (3% v/v for induction and 1% v/v for maintenance), a midline ventral incision in the anterior neck was made. HI pups underwent double ligation of the right common carotid artery with a 5.0 silk suture; Sham pups underwent surgery including incision and isolation of the right carotid artery without ligation. The entire surgery lasted ∼5-7min.

### S.2.7. P10 Seizure Scores

After carotid ligation or Sham surgery, the mice recovered for 60-80min on a heating pad with their littermates; temperatures of each individual pup were closely monitored and maintained between 36.0-37.9°C. All pups remained on heating pads until return to the dam. Recovery was followed by 45min of hypoxia exposure where oxygen concentration was decreased to 8% with balanced nitrogen in a controlled chamber (BioSpherix Ltd., Parish, NY, USA). The Sham mice remained the same amount of time in normoxia conditions (21% FiO_2_) in a similar chamber. During hypoxia as well as 10 min before and 10min after, mice were assessed on a modified Racine scale for seizure activity. Briefly, every 5min pups were assigned a score from 0 to 6 to reflect maximal seizure activity in that bin of time. Baseline behavior received a score of 0, behavioral arrest or immobility (1), head bobbing or rigid posture (2), unilateral limb clonus (3), bilateral limb clonus (4), exhibiting level 4 clonus for most of the 5min or level 4 for at least 30 seconds (5), and severe seizure activity with running or jumping (6). A cumulative seizure score was generated for each pup (ranging from 0-84). Across all experimental litters, the total time pups spent away from their dam was on average 120min. Within each litter, time away from the dam was calculated for HI pups and averaged; all Sham mice spent exactly this time away from the dam.

### S.2.8. Brain Collection and Tissue Preparation

Mice were killed at P10 or P13 via rapid decapitation using sterilized, surgical scissors in compliance with NIH guidelines for euthanasia of rodent neonates. Brains were then placed in room temperature (RT) 4% paraformaldehyde (PFA) and fixed for 72 hours. Following two PFA changes (1/day), brains were cryoprotected via placement in 30% sucrose in 0.1 M phosphate-buffered saline (PBS) at 4°C. Forty µm coronal sections were collected on a freezing microtome (Leica SM 2000 R) and stored in 1xPBS with 0.01% sodium azide at 4°C until processing for histological assessment. A subset of mice from the first cohort were not fixed, as detailed in the next section.

### S.2.9. Pale Surface Brain Lesion at P13: Presence and Size

For the presence of a pale lesion on the surface of the brain, each extracted brain underwent gross visual examination. If present, these pale brain surface lesions were always on the right hemisphere (due to right carotid artery ligation). For lesion size, if a pale surface lesion was present, a flexible ruler was placed on the surface of the brain to measure (in 0.5mm increments) the lesion at its greatest extent rostro-caudally and medio-laterally; calculated lesion area was the product of the length x width of the lesion.

### S.2.10 Immunohistochemistry (IHC) for Iba1

IHC was performed as previously described^57,58,69^ using an antibody against Iba1, which labels both brain-resident microglia and monocyte-derived microglia-like macrophages that infiltrate the brain from the peripheral blood. Briefly, 40µm brain sections were mounted onto charged slides (Thermo Fisher Scientific, #12-550-15) and left to dry for 1-2 h prior to IHC. Sections underwent antigen retrieval (0.01 M citric acid, pH 6.0, 100°C, 15min) followed by room temperature (RT) washes in 1xPBS; all subsequent steps also occurred at RT. Non-specific protein binding was blocked via incubation with 3% Normal Donkey Serum (NDS) and 0.3% TritonX-100 in 1xPBS for 60min. Following pretreatment and blocking steps, sections were incubated overnight with rabbit anti-Iba1 (1:5000; Wako Chemicals, #019-19741) in a carrier of 3% NDS and 0.3% Tween-20 in 1xPBS. Approximately eighteen hours later, sections were rinsed with 1xPBS prior to incubation with biotinylated donkey-anti-rabbit IgG (1:200; Jackson ImmunoResearch Laboratories, #711-065-152) in a carrier of 1.5% NDS in 1xPBS. After washing with 1xPBS, endogenous peroxidase activity was inhibited via incubation with 0.3% hydrogen peroxide in 1xPBS for 30min. Following an additional set of 1xPBS washes, slides were incubated with an avidin-biotin complex (ABC Elite, Vector Laboratories, #PK-6100) for 90min. Slides were washed with 1xPBS and immunoreactive cells were visualized via incubation with metal-enhanced diaminobenzidine (Thermo Fisher Scientific, #34065) for 11-12min. Slides were counterstained with Fast Red (Vector Laboratories, #H3403), dehydrated in a series of increasing ethanol concentrations and Citrosolv, and coverslipped with DPX (Thermo Fisher Scientific, #50-980-370).

### S.2.11. Quantification of Iba1-immunoreactive (+) cells

Results from Figures 1F, 2K, 5C used images from Iba1-immunostained (Iba1+) sections from P13 mice captured in bright-field light microscopy using an Olympus BX51 microscope with a 10x/0.3NA objective and an Olympus DP74 camera (widescreen aspect ratio, 16:10). The approach to quantify Iba1+ cells is described below.

#### S.2.11.1. Iba1+ pixel percentage in ipsilateral hippocampus after Sham or HI (Cohort 1)

Four 100x images of the ipsilateral (right) hippocampus per brain in Cohort 1 were obtained at -1.70, -2.30, -2.92mm relative to bregma, with two images at -2.92mm to capture the entire hippocampus at that bregma. Images were analyzed in Fiji. Raw brightfield images were converted to RGB stack and only the blue channel was analyzed because this provided the best signal-to-noise contrast for Iba1 signal.

Threshold was applied 0-141, and Fiji measurement reported the percentage of pixels positive for Iba1 immunoreactivity for each image. These were averaged across the four images per brain to obtain a single value representing the percent of Iba1+ pixels for the hippocampus of each mouse.

#### S.2.11.2. Iba1+ cell density in the left striatum, cortex, and hippocampus after Veh or PLX (Cohort 2)

For every mouse in Cohort 2, eight 100x images per brain were taken from the left hemisphere of five coronal sections ranging from +1.33 to -2.91mm from bregma; these sections included cortex, hippocampus, and striatum. Iba1+ cells were counted in on average 4 sections per brain from cortex, 3 sections per brain from hippocampus, 1 section per brain from striatum. Guidelines to ensure similar brain region representation were as follows: <u>Striatum</u>: the forceps minor of the corpus callosum was positioned in the top and lateral corner of the image. <u>Cortex</u>: the edge of the cingulum and corpus callosum were positioned at the bottom of the photomicrograph. <u>Hippocampus:</u> the dentate gyrus (DG) crest was placed in the lateral central screen such that the Hilus extended into the middle of the image. Images were examined in Fiji to measure the area of the region (striatum, cortex, hippocampus) in that section; the area of the brain region was converted to mm^2^ using the conversion 1pixel=9.19 x10^-7^mm^2^. Additionally, for Veh mice, we created a 400×400 pixel square (similar to Fig. 5A) in which Iba1+ cells were counted via Fiji’s multi-point tool. The number of Iba1+ cells in the square was divided by the area of the brain region (in mm^2^) to generate Iba1+ cell density of the given brain region. Iba1+ cell density was collected for each brain region (striatum, cortex, hippocampus) present in each photomicrograph. If multiple values were collected for a given brain region, they were averaged so that each mouse had one value for the striatum, cortex, and hippocampus. Within treatment, Iba1+ cell densities were similar between regions. Iba1+ cell densities from these regions were averaged such that each mouse was represented by one density value.

#### S.2.11.3. Iba1+ cells in the contralateral cortex after Veh or PLX and after Sham or HI (Cohort 4)

For every mouse in Cohort 4, one 100x image per brain was obtained at +1.21mm from bregma, maximizing the area within the image that contained the contralateral (left) cortex. In Fiji, the cortex was outlined using the polygon selection (to exclude corpus callosum and other subcortical regions). This area of the cortex was then measured and converted to mm^2^ using the conversion 1pixel=9.19 x10^-7^mm^2^. The number of Iba1+ cells in the cortex part of this image was counted using the multi-point tool for both Veh and PLX mice; however, they were exhaustively counted throughout the cortex image in PLX mice and counted within a 400×400 pixel square in Veh mice (see Fig 5A, B). For PLX mice, the calculated density of Iba1+ cells in the cortex was obtained by dividing the total number of Iba1+ cells in the cortex by the total area of the cortex for each 100x image. For Veh mice, the density of Iba1+ cells in the cortex was calculated by taking the number of Iba1+ cells in the square, multiplying by the total area of the cortex in the image, and dividing by the area of the 400×400 pixel square.

#### S.2.11.4. Iba1+ cells in the ipsilateral and contralateral hemisphere after Sham or HI in PLX mice (Cohort 4)

For every PLX mouse in Cohort 4, three sections (+1.21mm, -2.27mm, and -3.63mm from bregma) were scanned at 20x using a brightfield Aperio AT2 digital slide scanner from Leica Biosystems. Iba1+ cells were quantified using QuPath^59^. Using QuPath, a tight polygon was created around each hemisphere, yielding the area (in mm^2^) of the hemisphere. Within the tight polygon, Iba1+ cells were manually counted, resulting in the number of Iba1+ cells per mm^2^ for each hemisphere and section. An average of the Iba1+ cell density from each hemisphere across the three sections was used to determine Iba1+ cell density in the contralateral or ipsilateral hemisphere (cells/mm^2^), see Figures 5D-G.

### S.2.12. Cresyl Violet Staining and Lesion Scoring (Cohort 4)

Forty µm coronal brains sections in a 1:12 sampling ratio ranging from +1.41 to -3.63mm relative to bregma were mounted onto charged slides (Thermo Fisher Scientific, #12-550-15) and left to dry for 36 hours prior to staining with 0.1% cresyl violet acetate (CV). Cresyl violet stained sections from P13 mice used were analyzed in bright-field light microscopy using an Olympus BX51 microscope with both 10x/0.3NA and 20x/0.5NA objectives, and an Olympus DP74 camera (widescreen aspect ratio, 16:10). Lesion scoring for every section (range 7-11 sections per brain) was performed by two independent observers masked to experimental conditions in four hippocampal subregions (CA1, CA2, CA3, dentate gyrus), striatum, and cortex of all sections. The CA1 was defined as the most dorsal hippocampal subfield, formed by smaller, more dense pyramidal cell bodies than those observed in the CA2 and CA3. The CA2 was distinguished from the CA1 and CA3 by a widening of the pyramidal cell layer.^70^ Striatum was scored between bregma +1.41mm to -1.2mm because striatum was not present in sections posterior to this bregma. The cortex was scored in all sections from +1.41mm to -3.63mm. The left and right hemispheres were each scored independently. The entirety of each region within a section was scored on a scale of 0-3 with the following criteria: 0=no injury, 1=scattered shrunken neurons, 2=moderate cell loss with infarction in columnar distribution, 3=severe cell loss with cystic infarction.^60^ For each brain section, a score of 0-3 was given to each brain region analyzed. For each brain within each subregion, scores across bregma were averaged to yield average subregion lesion scores. Average subregion lesion scores and an average ipsilateral lesion score (average of all 6 subregions) from the two independent masked observers were averaged together to yield the final score graphed and analyzed; the average difference in lesion scores between observers was 1.6%. One observer scored all contralateral hemisphere sections; all scores were zero so the second observer scored only the ipsilateral hemisphere. To visualize lesion severity across groups in another way, the fraction of mice with average lesion score within each 0.5 increment bin was visually represented as parts of a whole.

### S.2.13. Statistical Analysis

Sample size estimation was based on prior results in comparable studies, assuming 80% power at a significance level of 0.05. Specific details on how many independent biological samples or mice were included in an experiment are given in the corresponding figure legends and in **Supplementary Table 1 (Supp. Table 1)**. All surviving mice were included in all statistical analyses except one Female-PLX-HI mouse. This mouse was an outlier in Iba1 cell counts (Grubbs test, G=3.331, *p<0.05) and its data were removed throughout; this mouse also had notable residue on its abdomen after PLX injections on both P9 and P10. These exclusion parameters were not determined *a priori*. Normality was assessed by the D’Agostino-Pearson omnibus test; homogeneity of variance was assessed by Spearman’s test. Parametric tests were preferentially used for analysis, including when data were close to but not normally distributed, given the robust nature of ANOVA testing per guidance from our statisticians. Data analyzed with parametric statistics are presented as mean ± standard error of the mean; data analyzed with non-parametric statistics are presented as median ± 95% confidence interval.

Three significant digits are used for statistical analysis reporting. Statistical analysis was performed using GraphPad Prism version 10.0.0 (GraphPad Software, USA) and R studio. Prism’s recommended *post hoc* tests were used when applicable. Statistical significance was defined at p<0.05 (asterisk). For effect size determination after ANOVA, the MOTE app^71^ or R studio were used to calculate partial omega squared (ω^2^) or generalized eta squared (ges; η^2^). These are reported with ≤0.01 small effect size, ≤0.06 medium effect size, and ≤0.13 large effect size ^72,73^, and 95% confidence interval (CI) is reported up to three significant digits. For effect size determination after non-parametric tests, partial eta squared is reported with ≤0.01 small effect size, ≤0.06 medium effect size, and ≤0.14 large effect size ^72,73^. For effect size determination of post-hoc analysis of multiple comparisons, Cohen’s d are reported for ANOVA results. Significant p values are bolded in **Table 1** and **Supp. Table 1** and reported rounded to the third decimal.

**Supplementary Table 1.**
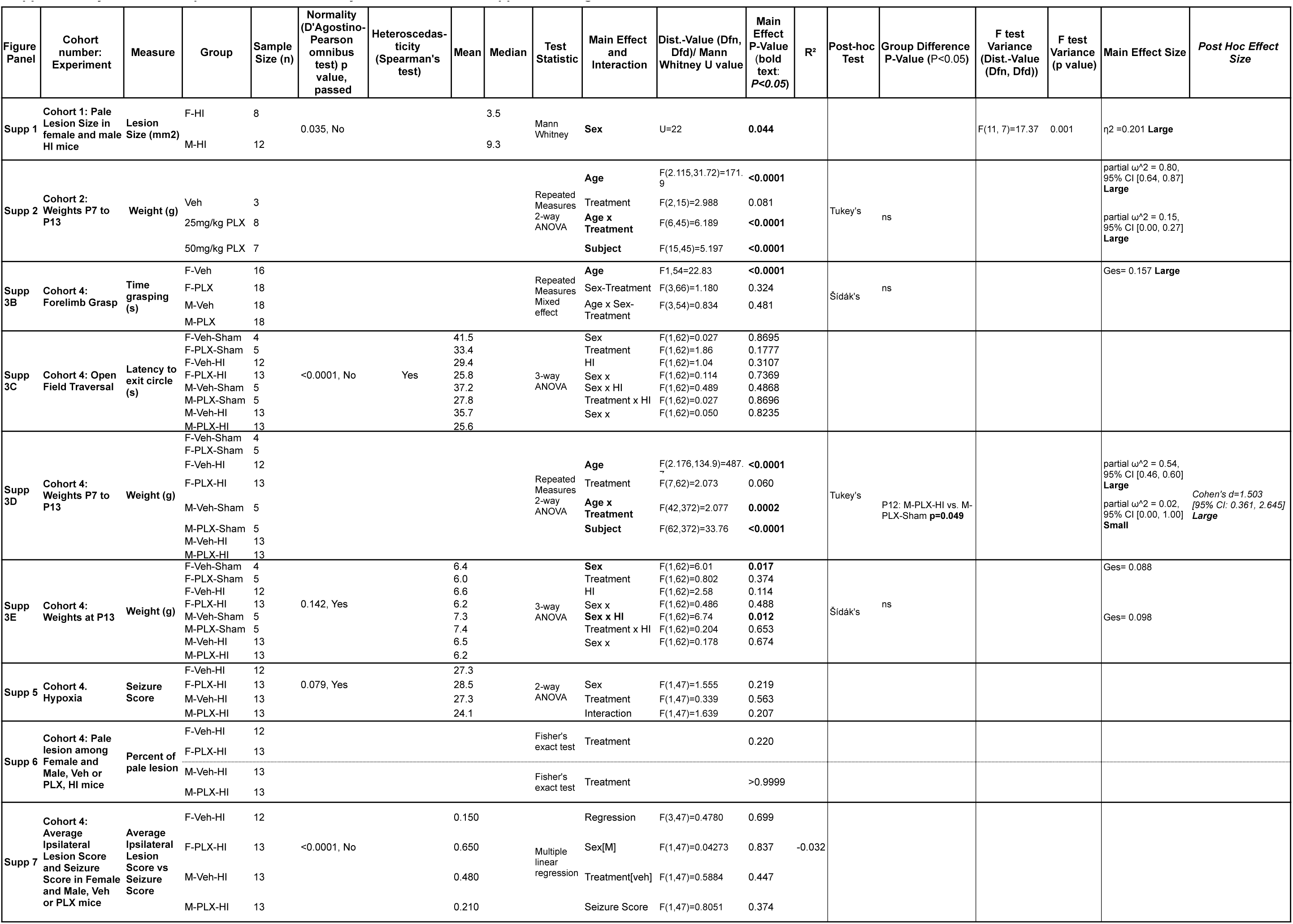
Sample Size, statistical analysis and results for supplemental figures.

## Notes

### Competing Interest Statement

The authors have declared no competing interest.

### Summary of Updates

Statisics table added for clarity. Figure 5 revised.

